# Ancestral evolution of oxidase activity in a class of (S)-nicotine and (S)-6-hydroxynicotine degrading flavoenzymes

**DOI:** 10.1101/2025.04.24.649827

**Authors:** Zhiyao Zhang, Parth R. Bandivadekar, Andrew J. Gaunt, Surl-Hee Ahn, Todd J. Barkman, Frederick Stull

**Affiliations:** Department of Chemistry, Western Michigan University, Kalamazoo, MI, USA; Department of Chemical Engineering, University of California, Davis, Davis, CA, USA; Department of Biological Sciences, Western Michigan University, Kalamazoo, MI, USA

**Keywords:** Flavin, oxidase, dehydrogenase, phylogenetics, molecular evolution, enzyme kinetics, molecular dynamics

## Abstract

Reduced flavin cofactors have the innate ability to reduce molecular oxygen to hydrogen peroxide. Flavoprotein oxidases turbocharge the reaction of their flavin cofactor with oxygen whereas flavoprotein dehydrogenases generally suppress it, yet our understanding of how these two enzyme classes control this reactivity remains incomplete. Here we used ancestral sequence reconstruction and biochemical characterization to retrace the evolution of oxidase activity in a lineage of nicotine/6-hydroxynicotine degrading enzymes of the flavoprotein amine oxidase superfamily. Our data suggest that the most ancient ancestor that gave rise to this lineage was a dehydrogenase, and that oxidase activity emerged later from within this group of dehydrogenases. We have identified the key amino acid replacements responsible for this emergence of oxidase activity, which, remarkably, span the entire protein structure. Molecular dynamics simulations indicate that this constellation of substitutions decreases the global dynamics of the protein in the evolution of oxidase function. This coincides with a dramatic restriction in the movement of a lysine residue in the active site, which more optimally positions it in front of the flavin to promote the reaction with O_2_. Our results demonstrate that sites distant from the flavin microenvironment can help control flavin-oxygen reactivity in flavoenzymes by modulating the conformational space and dynamics of the protein and catalytic residues in the active site.

## Introduction

Enzymes of the widespread flavoprotein amine oxidase (FAO) superfamily catalyze a diverse range of biologically important reactions and are present in all domains of life (1–3). FAOs all contain a flavin adenine dinucleotide (FAD) prosthetic group that they use to oxidize a carbon-nitrogen bond in an amine-containing substrate. The resulting reduced FAD hydroquinone (FADH_2_) then needs to be re-oxidized by a second substrate molecule to complete the catalytic cycle; most members of the FAO family are assumed to be oxidases that rapidly react with dioxygen (O_2_) to reoxidize their reduced flavin cofactor, forming H_2_O_2_ as a product (4–7). *Bona fide* flavin-dependent oxidases typically react with O_2_ with second order rate constants (k_ox_^O2^) on the order of 10^3^-10^6^ M^-1^s^-1^ (7–10). However, we recently showed that the FAO family member nicotine oxidoreductase (NicA2) is unusual in that it lacks typical oxidase activity, with the reaction between its FADH_2_ and O_2_ occurring with a very low k_ox_^O2^ of 28 M^-1^s^-1^ (11–14). Instead, NicA2 is a dehydrogenase, as its FADH_2_ is rapidly oxidized by a cytochrome c protein (CycN) that is encoded in an operon with *nicA2* in its native bacterium *Pseudomonas putida* S16 (11, 12). Genes for NicA2-like enzymes have only been found in bacteria of the Pseudomonadaceae family and all form an operon with a gene for cytochrome c, suggesting that all NicA2-like enzymes are dehydrogenases that react slowly with O_2_.

The closest homolog to NicA2 with a structure deposited in the PDB is the enzyme (S)-6-hydroxynicotine oxidase from *Shinella* sp. HZN7 (NctB), with the two enzymes sharing 43% sequence identity (15, 16). These two enzymes are structurally similar and oxidize very similar amine-containing substrates: (S)-nicotine for NicA2 and (S)-6-hydroxynicotine for NctB. However, while they utilize similar substrate molecules, they vastly differ in their rate of reaction with O_2_ (16, 17). NicA2 is a known dehydrogenase that reacts very slowly with O_2_ whereas NctB is a true oxidase that reacts rapidly with O_2_, with a k ^O2^ of 7.3 s^-1^ (16, 18). Moreover, phylogenetic analysis has shown that NicA2-like enzymes are closely related to NctB-like enzymes, with the two groups sharing a recent common ancestor that gave rise to the two sets of modern-day enzymes with highly divergent O_2_ reactivities (12, 19, 20). This highlights these enzymes as an interesting system for gaining new insight into how flavoenzymes evolve to gain or lose the ability to react rapidly with O_2_. Important questions to investigate about the evolution of these enzymes are: (i) was the last common ancestor of the NicA2-like and NctB-like enzymes an oxidase or a dehydrogenase? (ii) at what point during the evolution of these enzymes was the ability to react quickly with O_2_ gained or lost? (iii) which amino acid changes were required for the change in function, and how do these residues modulate the oxygen reactivity of the flavin prosthetic group?

We have used ancestral sequence reconstruction (ASR) to resurrect ancestral enzymes at relevant nodes to investigate the molecular basis for the evolution of O_2_ reactivity in the NicA2 and NctB containing lineage (21–25). Our data indicate that the last common ancestor between NicA2- and NctB-like enzymes was a dehydrogenase that reacted poorly with O_2_ and can efficiently use CycN as an oxidant. Potent oxidase activity then recently reemerged within a subset of (S)-6-hydroxynicotine-using enzymes, and many historical amino acid substitutions contributed to this substantial increase in oxygen reactivity. We found that these substitutions, which are both near and distant from the flavin, likely enhance the rate of reaction with O_2_ by dramatically decreasing the flexibility of a critical lysine residue, locking it in place proximal to the flavin C4a/N5 where the reaction with O_2_ occurs.

## Results

### Phylogeny of nicotine/6-hydroxynicotine degrading FAOs and ancestral sequence reconstruction

The top 500 sequences most similar to NctB from a protein BLAST search were used to construct a phylogenetic tree of NicA2- and NctB-like enzymes (Fig. S1A) (26, 27). As found previously, NicA2 and NctB are indeed related, with both enzymes present within a single clade that is phylogenetically isolated from other FAOs that function as oxidases. The NicA2 and NctB group enzymes diverged from a common ancestral enzyme at Node A early in the history of this clade, and since their divergence, they likely acquired a preference for different substrates: (S)-nicotine for NicA2 and (S)-6-hydroxynicotine for NctB (Fig. 1A and Fig. S1B) (2, 7, 18). NicA2 is a known cytochrome c-utilizing dehydrogenase, and an important genetic indicator of this enzymatic characteristic is the presence of a cytochrome c encoded within an operon with the *nicA2* gene (Fig. 1B). In contrast, enzymes of the NctB group have been confirmed to be oxidases and lack a cytochrome c gene encoded adjacent to *nctB* in the genome of origin (Fig. 1B) (18, 19). Directly sister to the NctB subclade is another subclade of enzymes from bacteria in the Sphingomonadaceae family, which we call the NctX group (Fig. 1A). These NctX enzymes are 55-60% identical to NctB enzymes and are found as part of a gene cluster in Sphingomonadaceae bacteria that contains the genes for catabolizing nicotine via the VPP pathway, indicating that NctX enzymes also likely act on (S)-6-hydroxynicotine as a substrate (15, 28, 29). Most notably, the genomes for NctX-containing organisms all encode a gene for cytochrome c that forms an operon with the *nctX* gene, strongly suggesting that NctX enzymes, like NicA2, are cytochrome c using dehydrogenases (Fig. 1B). Unfortunately, we were unable to produce a modern day NctX in soluble form through heterologous expression in *E. coli* for biochemical characterization, which may be due to the fact that NctX enzymes are predicted to be membrane anchored lipoproteins (30). However, our characterization of ASR constructs described below is consistent with modern day NctX enzymes being cytochrome c-using dehydrogenases. Also noteworthy for the purposes of this study are the two FAO protein sequences from an unspecified Burkholderiaceae bacterium and *Pseudohalioglobus lutimaris* in Figure 1A, as our phylogenetic analysis indicates that these two modern day sequences diverged earlier along the lineage that gave rise to the NctB and NctX subclades. Both of these FAOs have a cytochrome c gene forming an operon with their *fao* gene, suggesting that these two modern-day enzymes also function as dehydrogenases.

**Fig. 1.**
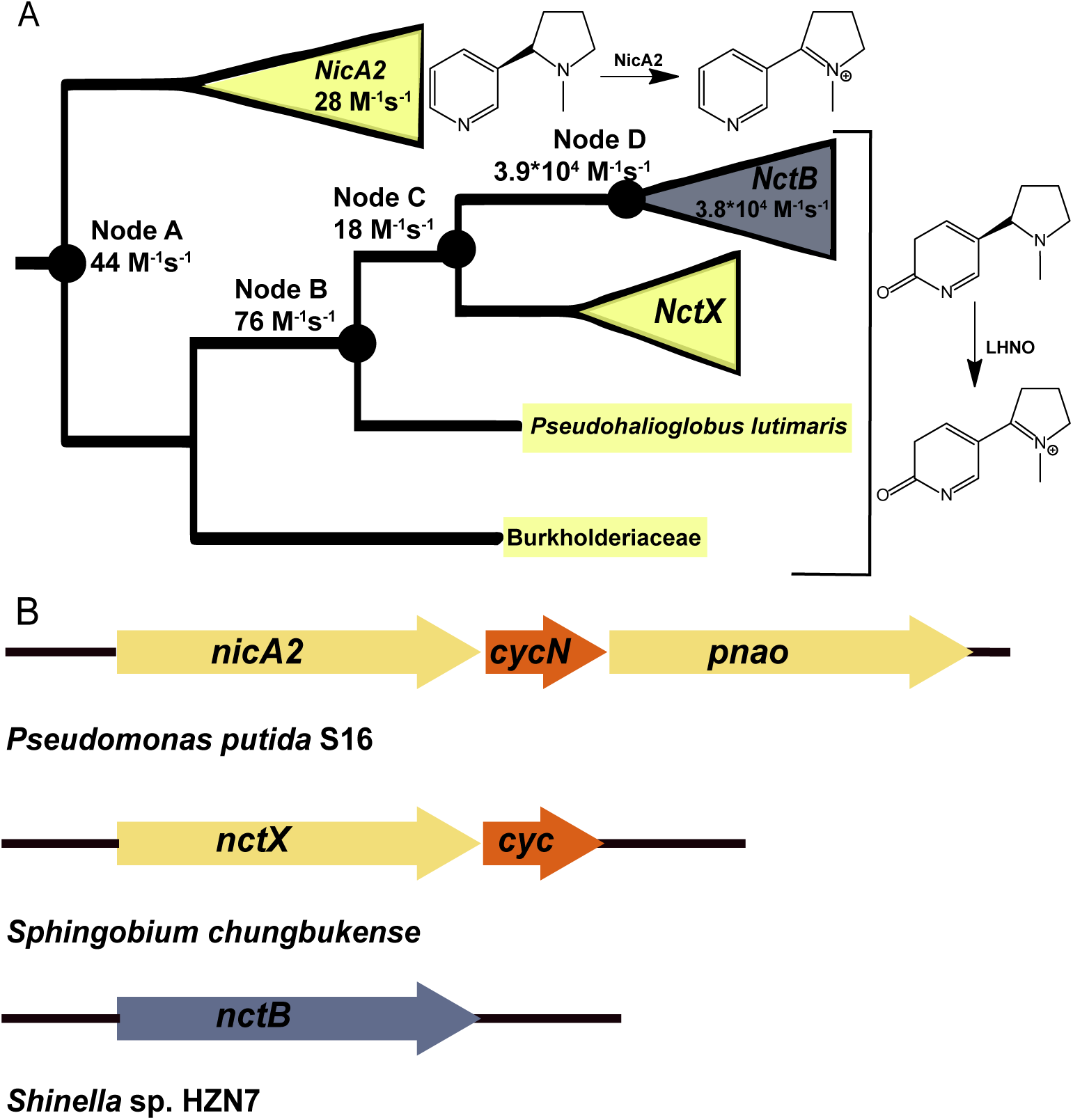
Nicotine/6OHN degrading FAOs are closely related and include both dehydrogenases and oxidases. (A) Collapsed phylogeny of nicotine/6OHN degrading FAOs with targeted nodes for ASR represented by filled circles. Clades are colored based on the presence (yellow) or absence (blue) of genome-adjacent cytochrome c. The second order rate constant for reaction with O_2_ (k_ox_^O2^) is listed for the modern-day enzymes and ancestral nodes characterized in this study. (B) Example genome arrangements for FAOs and neighboring cytochrome c genes or lack thereof.

The most parsimonious hypothesis from the aforementioned presence of adjacent cytochrome c genes is that the most ancient ancestor that gave rise to the NicA2 and NctB/NctX lineages (Node A) was a cytochrome c-using dehydrogenase. If true, then the oxidase activity of NctB evolved from within this clade of dehydrogenases, most likely at either Node C or Node D. To test this hypothesis and identify the branch in ancestral history where oxygen reactivity changed, we selected four nodes from the phylogenetic tree and generated ASR constructs of them for biochemical characterization (Fig. 1A). Node A is the most ancient ancestor for all of the NctX, NctB and NicA2 groups. Node B is the ancestral node where the FAO from *Pseudohalioglobus lutimaris* diverged from the lineage that led to the NctX/NctB groups. Node C is the ancestor that gave rise to the NctX and NctB groups, and Node D is the immediate ancestor to the NctB group enzymes. All four ancestral nodes had high branch support (>0.99) (Fig. S1B) and the average posterior probability for the ancestral sequence estimates ranged from 0.85-0.92. The genes corresponding to these ancestral proteins were synthesized and cloned into pET29b as his-tagged constructs and the proteins were expressed and purified. All ASR constructs purified with FAD bound, indicating that they were properly folded.

### Substrate specificity of ancestral FAOs

Modern-day NicA2 enzymes use (S)-nicotine as their amine-containing substrate, whereas modern day NctB and NctX enzymes use (S)-6-hydroxynicotine (6OHN) as their substrate, indicating that the switch to using 6OHN occurred somewhere along the lineage that gave rise to NctB/NctX enzymes. We therefore investigated the kinetics of reaction with nicotine or 6OHN for the ancestral enzymes at Nodes A through D using anaerobic stopped-flow experiments to determine when the switch in substrate preference occurred (Fig. 2). For reference, we also measured the kinetics of reaction with nicotine or 6OHN for modern-day NctB from *Shinella* sp. HZN7 (Fig. 2A). As purified, all of the ASR enzymes and NctB have typical spectra of oxidized FAD. Following the addition of nicotine or 6OHN, the oxidized FAD is reduced to FADH_2_ with a corresponding loss of absorbance at 450 nm, and this signal was used to monitor the kinetics of the reaction with nicotine or 6OHN in the absence of O_2_. NctB, Node B, Node C, and Node D can be reduced by nicotine or 6OHN, with the reaction traces generally occurring in two kinetic phases with comparable amplitudes (Fig. S2). We previously observed similar biphasic behavior in NicA2’s reaction with nicotine, which we attributed to the two active sites of the NicA2 homodimer reacting with different kinetics (11). Exceptions to this biphasic behavior were observed with the reaction of Node B with nicotine and Node C with nicotine, with only a single reaction phase observed in both cases. Notably, the reaction with 6OHN was completed within 1–10 seconds for Nodes B, C, and D, whereas the reaction with nicotine took more than 100 seconds to complete, indicating a strong preference for 6OHN as a substrate for these three ancestral enzymes (Fig. 2C, 2D & 2E). Consistent with this, the rate constants for flavin reduction (k_red_) are >80-fold lower with nicotine than with 6OHN for Nodes B, C, and D (Table S1). Similar reaction times are observed with modern day NctB from *Shinella* sp. HZN7 (Fig. 2A), supporting the notion that 6OHN is the preferred substrate for the Nodes B, C, and D enzymes. In contrast, Node A reacted poorly with both nicotine and 6OHN, taking more than 800 seconds to achieve only partial FAD reduction with both potential substrates (Fig. 2B). This observation may indicate that the Node A enzyme used a substrate other than nicotine or 6OHN, and that preference for 6OHN evolved between Node A and B and then persisted along this branch to modern day NctB and NctX enzymes.

**Fig. 2.**
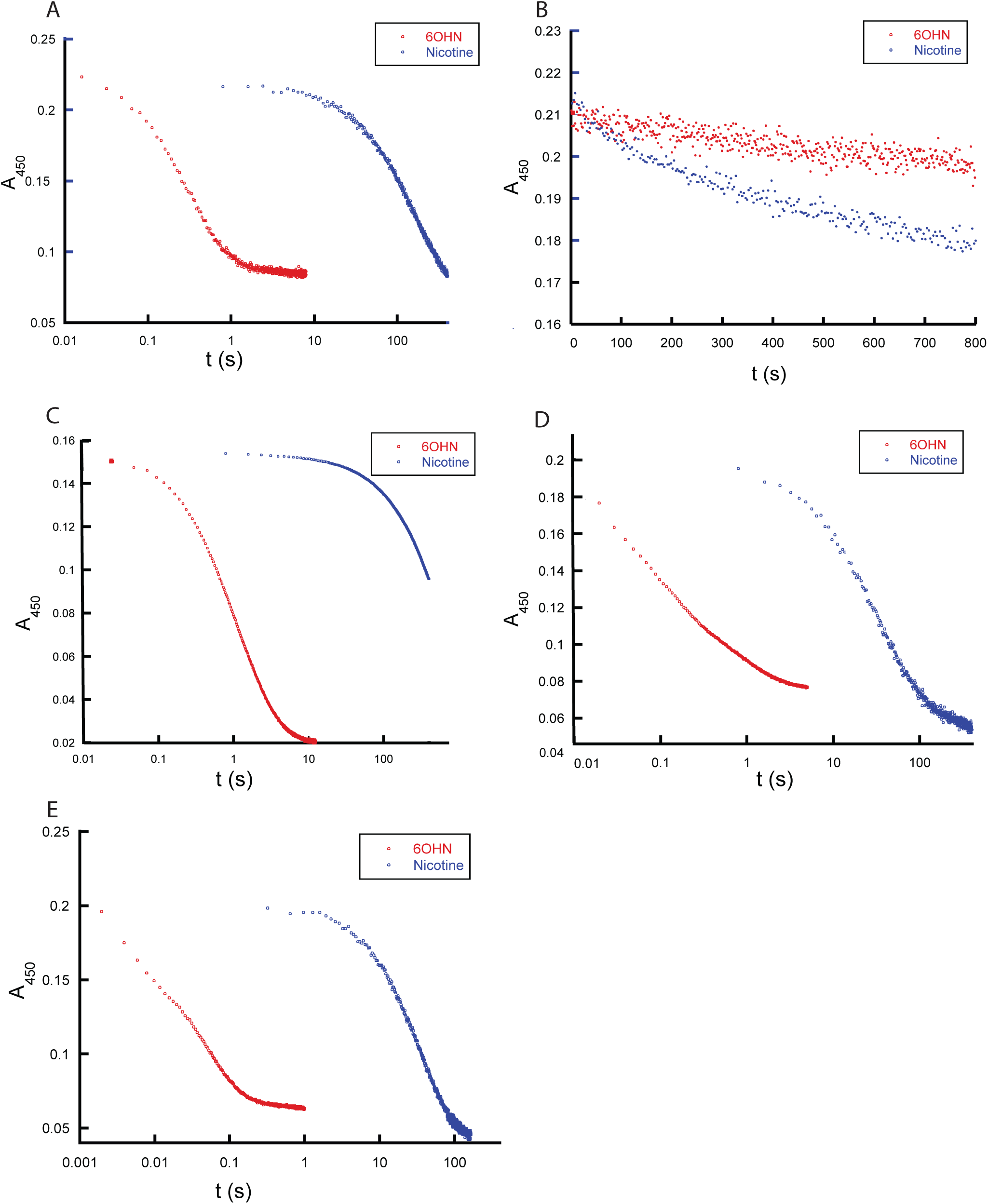
Kinetics of flavin reduction by (S)-nicotine or (S)-6-hydroxynicotine. (A), (B), (C), (D), and (E) Data for NctB, Node A, Node B, Node C and Node D, respectively. A stopped-flow reaction trace of each oxidized flavoenzyme with 200 µM of (S)-6-hydroxynicotine (blue) or (S)-nicotine (red). Note the logarithmic time scale. Reaction traces at other substrate concentrations and observed rate constant data can be found in Fig. S2 and Table S1.

### Oxidase activity recently reemerged

After investigating their reductive half reactions, we then tested each of the reduced ancestral enzymes and modern day NctB for their ability to react with O_2_ in stopped-flow experiments by monitoring flavin oxidation at 450 nm (Fig. 3A-3E). Fitting reaction traces to an exponential function provided an observed rate constant (k_obs_) value for each trace, and subsequent fitting of k_obs_ against O_2_ concentration to a line provided the second order rate constant for flavin oxidation by O_2_ (k_ox_^O2^). As shown in Fig. 1A, Fig. 3F and Table 1, Node A, Node B and Node C reacted very slowly with O_2_, producing k_ox_^O2^ values of 44±1 M^-1^s^-1^, 76±1 M^-1^s^-1^, and 18±1 M^-1^s^-1^, respectively, all of which are >2 orders of magnitude lower than typical oxidases (10^3^∼10^6^ M^-1^s^-1^) and very close to the value measured for modern-day NicA2 which is a *bona fide* dehydrogenase (11). In contrast, Node D reacted very quickly with O_2_, showing typical oxidase reactivity with a k_ox_^O2^ of 3.9±0.1*10^4^ M^-1^s^-1^, similar to the value of modern day NctB, which has k_ox_^O2^ of 3.8±0.1*10^4^ M^-1^s^-1^ (Fig. 3F & Table 1). These data indicate that the oxidase activity in NctB emerged from ancestral enzymes that were likely dehydrogenases, and that oxidase activity evolved during the divergence between Node C and Node D (Fig. 1). The data also indicate that the NctX group are likely dehydrogenases since their direct ancestor, Node C, reacts poorly with O_2_, and *nctX* forms an operon with a gene for cytochrome c in NctX-containing organisms.

**Fig. 3.**
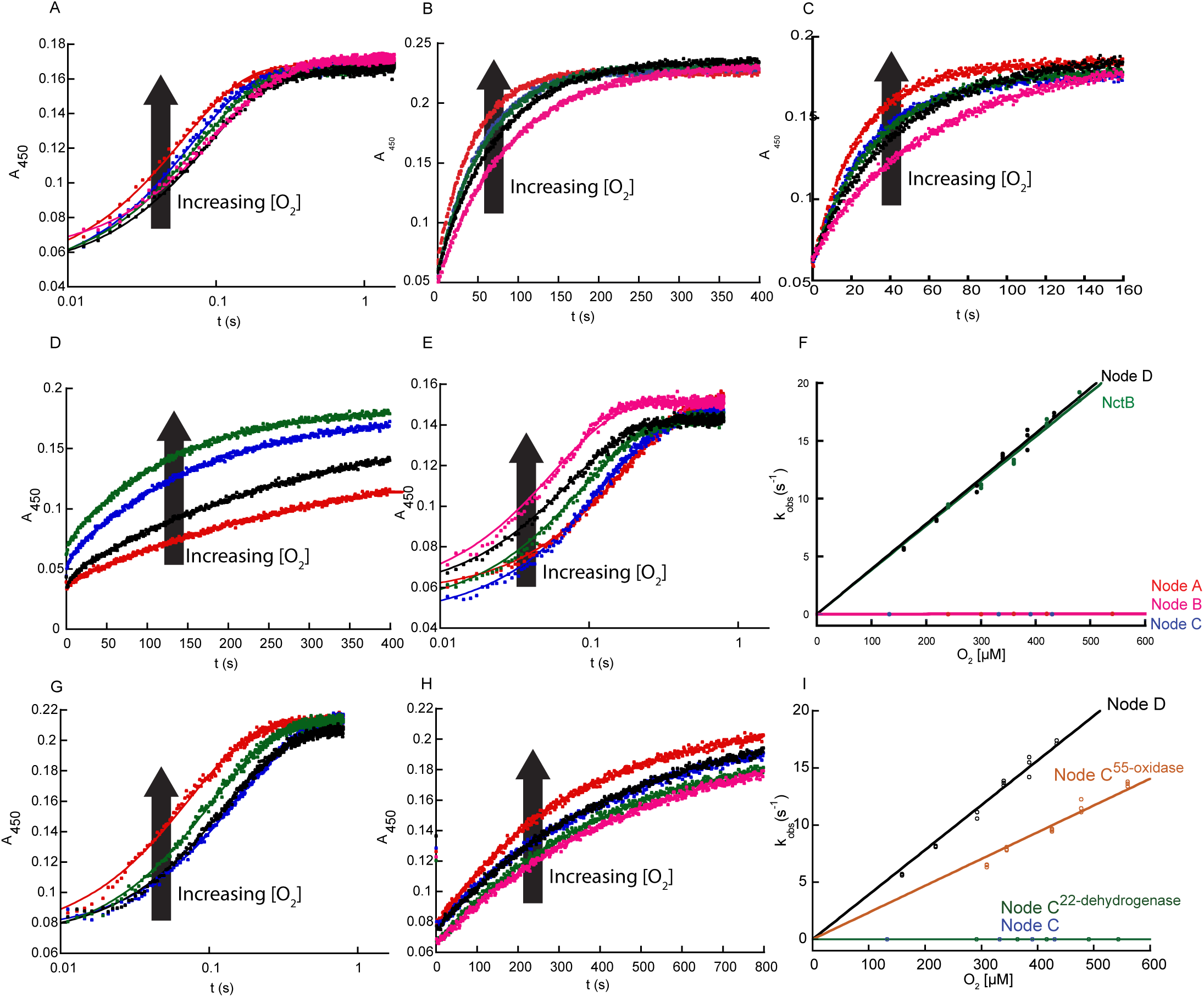
Kinetics of flavin oxidation by O_2_. (A), (B), (C), (D), and (E), Stopped-flow absorbance traces at 450 nm for the oxidation of reduced NctB, Node A, Node B, Node C, and Node D, respectively, by various concentrations of O_2_. (F) Plots of k_obs_ values against O_2_ concentration for the reaction of various reduced ancestral FAOs with O_2_. (G) and (H) Stopped-flow absorbance traces at 450 nm for the oxidation of reduced Node B^55-oxidase^ and Node B^22-dehydrogenase^, respectively, by O_2_. (I) Plots of k_obs_ values against O_2_ concentration for the reaction of Node C^55-oxidase^ and NodeC^22-dehydrogenase^ with O_2_. In panels F and I, the slope obtained from linear fitting gave the k_ox_^O2^ values for the FAOs reported in Table 1.

**Table 1.** Rate constants for reaction <u>with O_2_</u>.

| Enzyme | $k_{\text{ox}}^{\text{O}_2} (\text{M}^{-1}\text{s}^{-1})^{\text{a}}$ |
| --- | --- |
| Node A | 44±1 |
| Node B | 76±1 |
| Node C | 18±1 |
| Node C<sup>22</sup>-dehydrogenase | 7.0±0.5 |
| Node C Thr300Glu | 170±10 |
| Node C<sup>5</sup> | 160±10 |
| Node C<sup>13</sup> | 3.7±0.3*10<sup>3</sup> |
| Node C<sup>21</sup> | 1.9±0.2*10<sup>4</sup> |
| Node C<sup>55-oxidase</sup> | 2.3±0.1*10<sup>4</sup> |
| Node D | 3.9±0.1*10<sup>4</sup> |
| NctB | 3.8±0.1*10<sup>4</sup> |

<sup>a</sup>Error values denote standard errors.

### Cytochrome c may be the preferred oxidant for ancestral Nodes A–C

The low rate constant of reaction with O_2_ for Nodes A, B, and C indicates that these ancestral FAOs likely use some electron acceptor other than oxygen as their preferred oxidant. Since NicA2 is a known cytochrome c-using dehydrogenase, and NctX group enzymes presumably also use a cytochrome c (due to the presence of the cyt c gene adjacent to all *nctX* genes), a cytochrome c is the most obvious candidate oxidant used by Nodes A–C. The cytochrome c from *P. putida* S16 (CycN) used by NicA2 was used to investigate this hypothesis. Anaerobic stopped-flow experiments were conducted to measure the kinetics of electron transfer from reduced ancestral enzymes to oxidized CycN (CycN_ox_), with the reaction monitored at 550 nm corresponding to cytochrome c reduction (Fig. 4). The reaction between Node A and CycN was rapid and completed within one second (Fig. 4A). Kinetic traces for this reaction required a linear combination of three exponentials to fit, with the first two kinetic phases having similar amplitudes and comprising ∼85% of the total signal change. The observed rate constant for the first two phases increased linearly with CycN concentration and were attributed to two stepwise one electron transfers from reduced flavin to two molecules of CycN, both rate-limited by binding to CycN, similar to that observed for NicA2 (Fig. 4D) (11, 31). The origin of the third, small phase is unclear but may be due to a small population of damaged enzyme. Linear fitting of the plots of k_obs_ against CycN concentration yielded second order rate constants (k_ox_^CycN^) of 4.5±0.2 *10^4^ M^-1^s^-1^ and 8.5±0.5 *10^3^ M^-1^s^-1^ for the first and second phase, respectively (Table 1). These values are comparable to those measured previously for the reaction of NicA2 with CycN, in agreement with Node A being a cytochrome c using dehydrogenase (11). Node B and Node C can also react with CycN, but the reactions are considerably slower, taking roughly 80 seconds to complete (Fig. 4B and 4C). Kinetic traces for both enzymes fit best to a linear combination of two exponentials, and k_obs_ increased linearly with CycN in all cases (Fig. 4E and 4F). The second order rate constants for the two kinetic events for Node B were 1.2±0.1*10^3^ M^-1^s^-1^ and 390±10 M^-1^s^-1^ and for Node C were 1.3±0.3*10^3^ M^-1^s^-1^ and 330±10 M^-1^s^-1^ (Table 1). These values are notably >10-fold lower than the corresponding ones for the reaction between Node A and CycN. We have previously observed that modern day FAO dehydrogenases NicA2 and Pnao display specificity for the native cytochrome c that they use, suggesting that the sequences of these putative ancestral FAO dehydrogenases may have been optimized to react with specific ancestral cytochrome c partners with a different sequence from CycN (11, 32). Node A has a more similar sequence to NicA2 (79% identity) than Node B or Node C (54% and 51% identity), and this may explain why Node A reacts more efficiently with CycN than Node B or Node C. Regardless, the first k_ox_^CycN^ for Node B and Node C is >15-fold larger than the corresponding k_ox_^O2^ values, which are very low for both enzymes, in agreement with Node B and Node C being cytochrome c using dehydrogenases. We also evaluated the ability of reduced Node D and NctB to react with CycN. Reaction traces for both of them were very slow and showed only a minor increase in absorbance at 550 nm relative to that seen with Nodes A–C (Fig. 4G and 4H). This indicates that only a small percentage of the reduced Node D and NctB enzymes were oxidized by CycN, and that CycN is a poor oxidant for Node D and NctB (Fig. 4G and 4H). These data, combined with the analysis of oxidase activity above, suggests that ancestral Nodes A–C were cytochrome c using dehydrogenases with low reactivity towards O_2_; this dehydrogenase function was lost during the divergence between Node C and Node D, with an associated dramatic increase in the rate of reaction with O_2_.

**Fig. 4.**
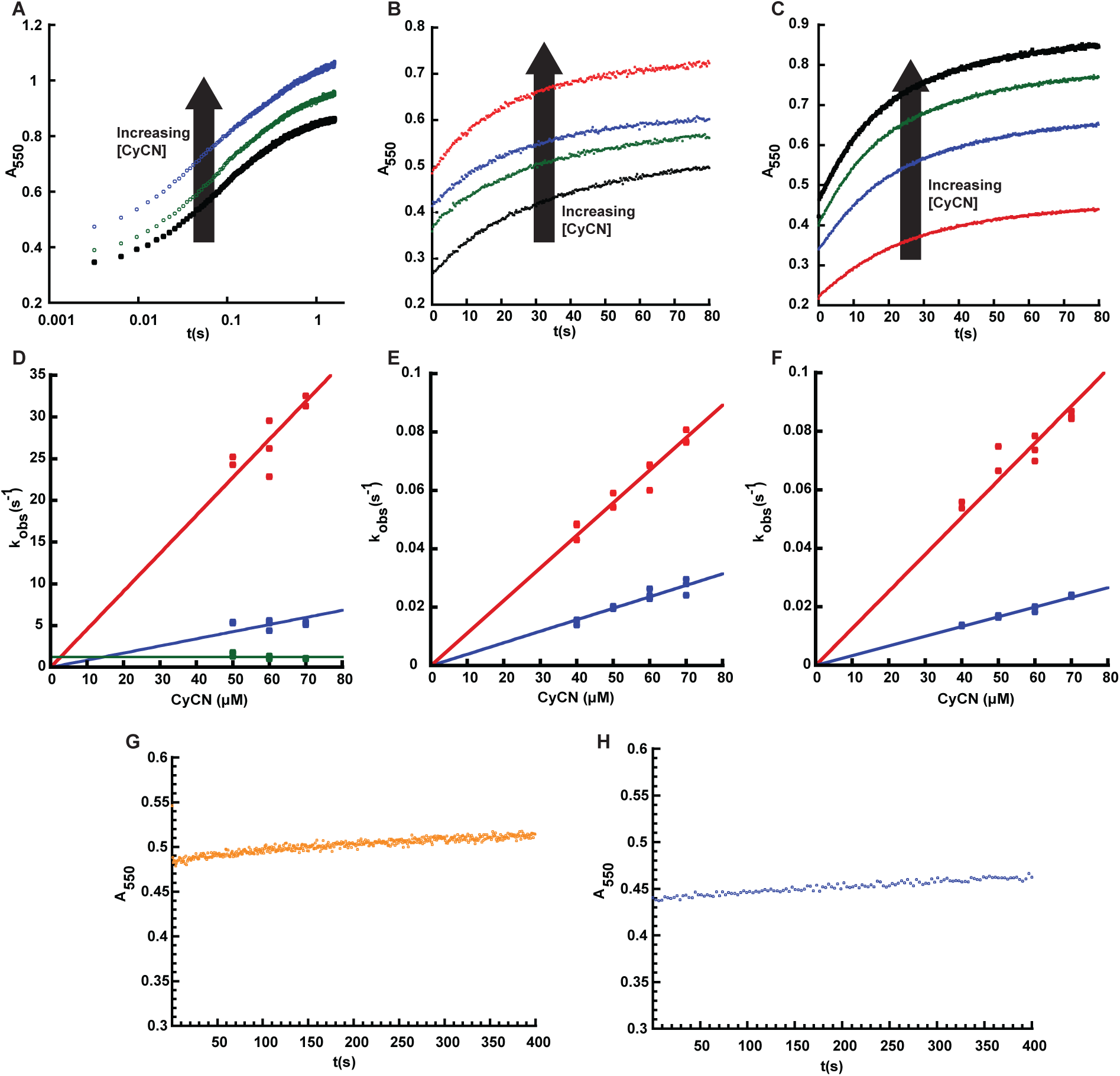
Kinetics of flavin oxidation by CycN. (A), (B) and (C) Absorbance traces at 550 nm for the stopped flow reaction between reduced Node A, Node B and Node C, respectively, and variable concentrations of oxidized CycN. Note the logarithmic time scale in panel A. (D), (E) and (F) Plots of k_obs_ values against CycN concentration for the reaction of Node A, Node B and Node C with CycN, respectively. The slope obtained from linear fitting gives the k_ox_^CycN^ values for the ancestral nodes. The values can be found in Table 1. (G) and (H) Absorbance traces at 550 nm for the stopped flow reaction between reduced Node D and NctB respectively, with 50 µM of CycN.

### Identifying key residues for oxidase activity

Because the emergence of potent oxidase activity in the NctB lineage occurred between Nodes C and D (Fig. 1A), we sought to identify potential amino acid residue positions that also changed during this divergence, since they must be responsible for the evolution of this enzymatic property. Node C and Node D share 82% sequence identity, with 77 of the 431 amino acid positions varying between them (Fig. S3). To minimize the number of amino acid replacements for downstream biochemical analysis, we compared the amino acid identities at these 77 positions with all 31 extant sequences shown in Fig. 1A, and the result is shown in Fig. S4. We hypothesized that if Node D had a particular residue at a given sequence position that was not found in any of the dehydrogenases of the alignment, then it could be important for oxidase activity. Using this criterion, we identified 55 sites that uniquely changed between Nodes C and D. However, for the other 22 sites that were inferred to change between Nodes C and D, the Node D enzyme (and/or its descendant NctB oxidases) possessed a residue that some of the dehydrogenases also contained. Because some of the proteins in the dehydrogenase clades also shared the Node D residue, we reasoned that those sites would likely not be important for the evolution of oxidase activity. To evaluate the potential functional importance of the putative oxidase sites, we simultaneously replaced the 55 residues in Node C with the Node D residues found at homologous positions (Node C^55-oxidase^). As shown in Fig. 3G & 3I and Table 1, the resulting k_ox_^O2^ value for Node C^55-oxidase^ is 2.3±0.1*10^4^ M^-1^s^-1^, comparable to the value of Node D, suggesting that all or a subset of these residues were involved in the evolution of oxidase activity from an ancestral dehydrogenase. In contrast, when we introduced the 22 positions that were hypothesized not to be important for the evolution of oxidase activity between Nodes C and D into the Node C ancestral enzyme, Node C^22-dehydrogenase^, the resulting k_ox_^O2^ value was 7.0±0.5 M^-1^s^-1^ (Fig. 3H & 3I). This value is comparable to Node C and other *bona fide* dehydrogenases, thereby indicating that these sites are probably not crucial for the evolution of oxidase activity in this lineage.

We next used site-directed mutagenesis to individually mutate residues in Node D towards those found in Node C, to generate 55 “reverse” (D➔C) mutants that were subsequently purified and characterized. We chose to analyze reverse mutants instead of “forward” (C➔D) mutants because we hypothesized that the gain in oxidase activity between Node C and Node D might require "permissive" epistatic amino acid replacements; if true, many of the 55 individual "forward" mutations would likely be uninformative alone (33). All 55 single mutants were screened for oxidase activity in steady state assays using an oxygen probe to monitor the rate of oxygen consumption with 6OHN as the substrate (Fig. S5). Of the 55 single mutants tested, 21 of them showed a 25% or greater reduction in activity relative to Node D (Fig. 5). Five single mutants reduced activity by more than 60%, and three of these (Glu300Thr, Met100Val and Ala90Val) dramatically reduced the oxygen consumption rate to a level similar to Node C (Fig. 5).

**Fig. 5.**
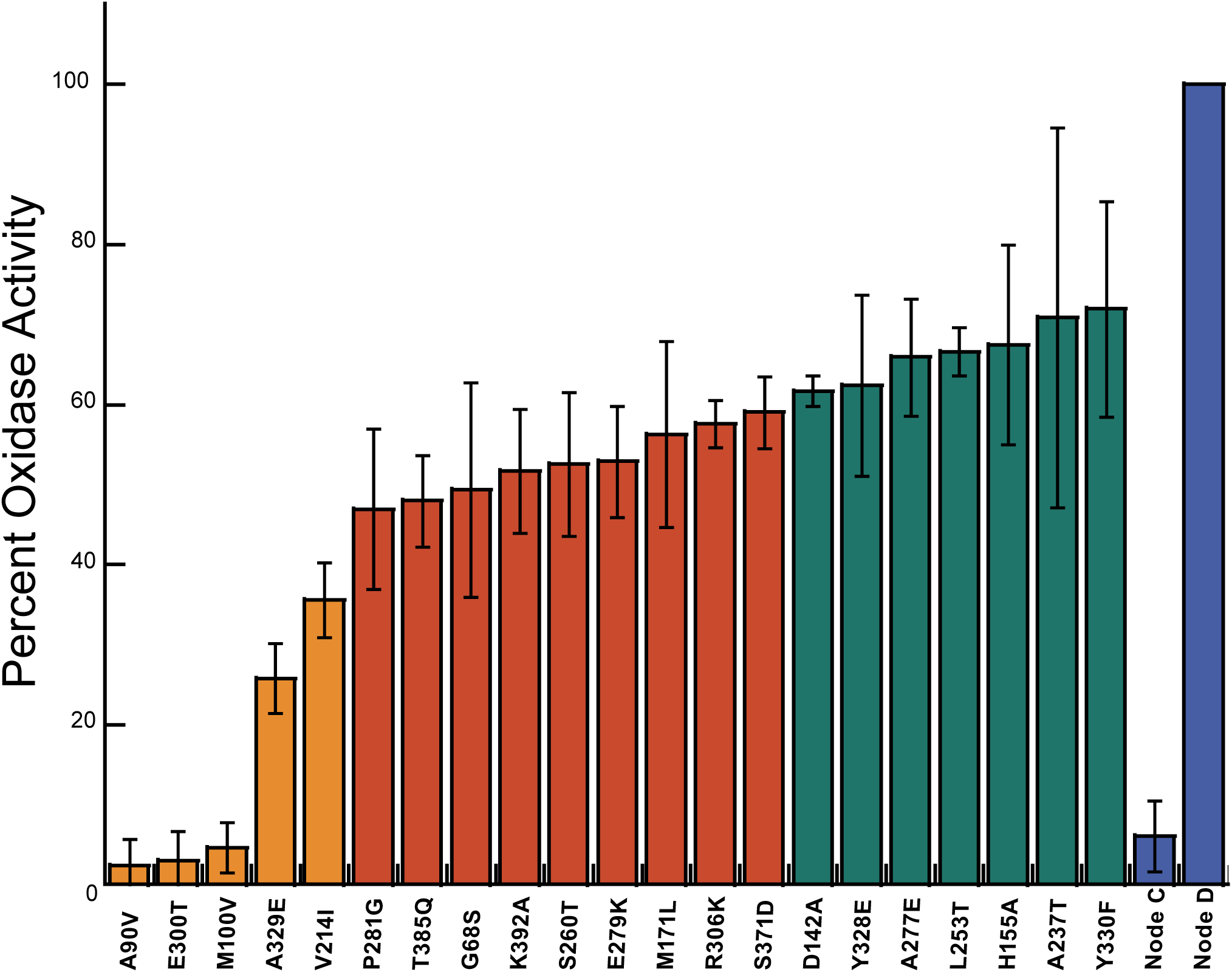
Relative oxidase activity of D➔C single mutants using 6OHN as a substrate. Relative oxidase activity was calculated using equation 5. Residue replacements with >60%, 60-40% and 40-25% decrease in relative reaction rate are colored with orange, red, and green, respectively. Note that numbers represent the residue position in Node D and replacement information indicates the amino acid in Node D that is replaced with the residue present in Node C for each D➔C variant. Data for additional variants can be found in Fig. S5. Error bars represent standard deviations.

### Glu300 is critical for rapid FADH_2_ oxidation by O_2_ in Node D

We selected the Glu300Thr, Met100Val, and Ala90Val D➔C single mutant enzymes for in-depth analysis by stopped-flow since these three substitutions reduced the O_2_ consumption rate in steady state assays to that observed for Node C. We first analyzed the kinetics of the reductive half-reaction for each mutant by anaerobically mixing enzyme containing oxidized flavin with 6OHN in stopped-flow experiments, using the change in absorbance that accompanies flavin reduction as a readout (Fig. S6A-F). All three mutants were capable of reacting with 6OHN, but only the Glu300Thr D➔C mutant did so with rate constant(s) comparable to the Node D (Table 1). Both the Ala90Val and Met100Val D➔C variants showed prolonged reaction times with 6OHN, taking more than 100 seconds to complete (Fig. 6A). Fitted data for Ala90Val and Met100Val reacting with 6OHN revealed 100- and 10-fold smaller k_red_ values, respectively, for both phases, indicating a compromised ability of these mutants to react with 6OHN (Table S1). This result is surprising given that the Node C and Node D enzymes used to derive these D➔C mutants both react rapidly with 6OHN, and positions homologous to Ala90Val and Met100Val are far from the substrate binding site in the structures of NicA2 and NctB. We then tested the Ala90Val, Met100Val, and Glu300Thr mutants’ ability to react with O_2_ in stopped-flow experiments to measure k_ox_^O2^ (Fig. S6G-I). The Met100Val variant had a k_ox_^O2^ of 2.8 ± 0.1 *10^4^ M^-1^s^-1^, a modest 1.4-fold decrease relative to the parent Node D (Fig. 6B and Table S1). In contrast, the Ala90Val and Glu300Thr variants had k_ox_^O2^ values of 7.0±0.1*10^3^ M^-1^s^-1^ and 1.1±0.2*10^3^ M^-1^s^-1^, indicating that reactivity with O_2_ is indeed significantly compromised by these mutations (Fig. 6B and Table S1). Comparison of k_red_ and k_ox_^O2^ with the 6OHN and O_2_ concentrations used in our activity assays indicates that the dramatic decline in steady state activity observed for the Ala90Val and Met100Val mutants in Fig. 5 is likely due to an impaired ability to react with 6OHN whereas the Glu300Thr mutant is rate-limited by its reaction with O_2_.

**Fig. 6.**
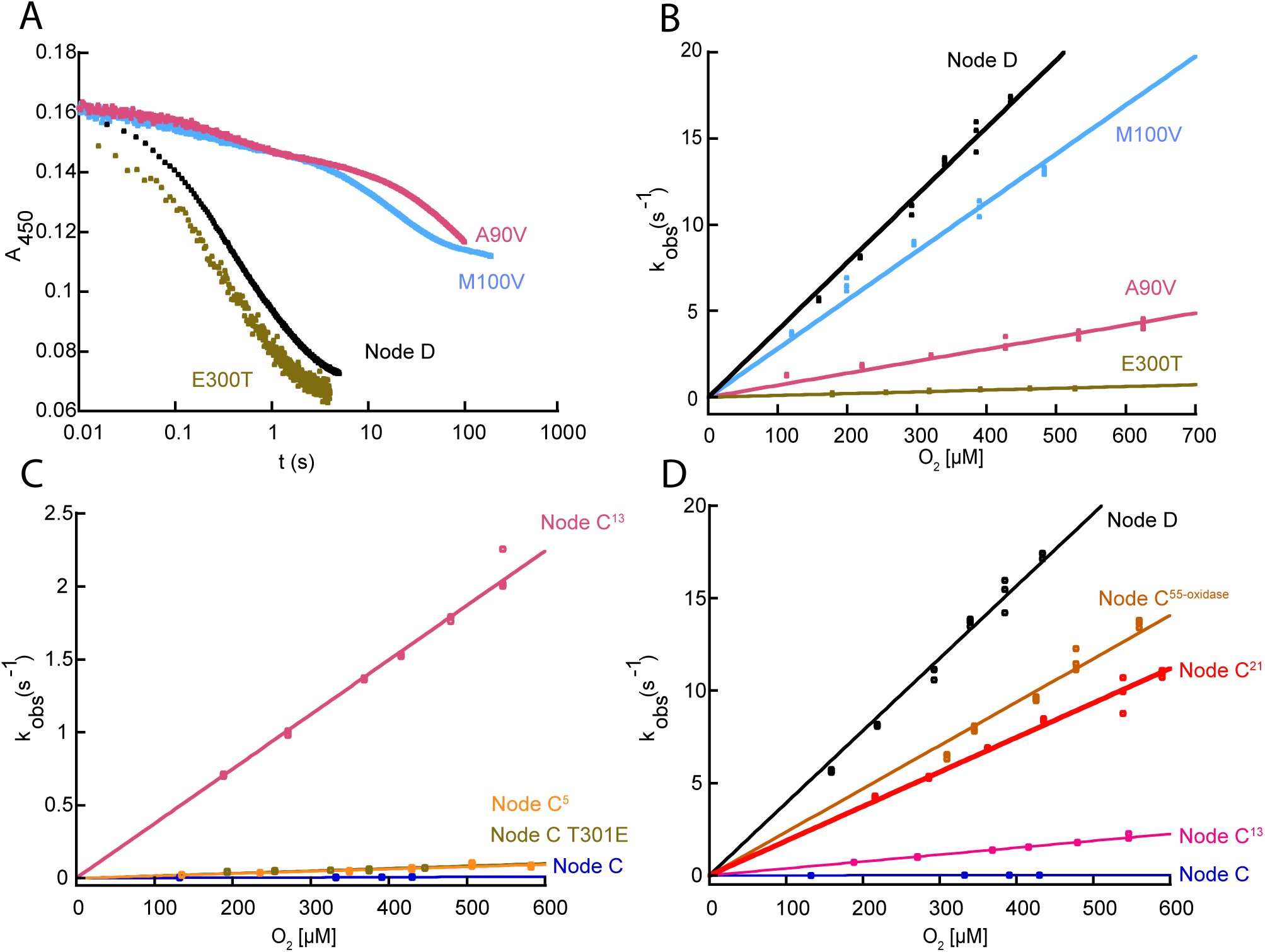
Stopped-flow kinetics for D➔C single mutants and C➔D variants. **(A)**, Stopped-flow absorbance traces for different D➔C single mutants reacting with 2 mM 6OHN to show differences in 6OHN reactivity. Note the logarithmic time scale. **(B)** Plots of k_obs_ values against O_2_ concentration for the reaction of different reduced D➔C single mutants with O_2_. Data for Node D is included for reference. **(C)** and **(D)** Plots of k_obs_ values against O_2_ concentration for the reactions of different reduced C➔D variants with O_2_. Data for Node C and Node D are included for reference.

### Converting Node C into an oxidase

After identifying residues that contribute to oxidase activity from the “reverse” mutant screening of Node D➔C variants (Fig. 5), our next objective was to attempt to convert the Node C ancestral dehydrogenase into an oxidase. This was achieved by introducing different nested subsets of the functionally important residues identified by the Node D➔C mutagenesis screening into Node C (making Node C➔D variants). Four Node C➔D variants were constructed, purified and k_ox_^O2^ was determined for each in stopped-flow experiments: Node C Thr300Glu contained only the single substitution of largest effect on oxygen reactivity from our Node D➔C mutant screening and stopped-flow experiments; Node C^5^ contained the five mutations that produced a >60% decline in activity from our D➔C screening (Val90Ala, Val100Met, Ile214Val, Thr300Glu and Glu329Ala); Node C^13^ contained the thirteen mutations that produced a >40% decline in activity; Node C^21^ contained the twenty-one mutations that produced a >25% decline in activity (see Fig. 5 for amino acid replacements). Importantly, stopped-flow experiments to directly measure k_ox_^O2^ ignore any effect that the mutations may have on the reaction with 6OHN. Stopped-flow reaction traces for these experiments are shown in Fig. S7A-D. Node C Thr300Glu displayed a modest 9-fold increase in k_ox_^O2^. While Node C exhibited a k_ox_^O2^ of 18±1 M^-1^s^-1^ the k_ox_^O2^ for Node C Thr300Glu was 170±10 M^-1^s^-1^, which confirms that the Thr300Glu substitution contributes to O_2_ reactivity (Table 1 and Fig. 6C), though, on its own, is insufficient to achieve the potent oxidase activity of Node D. Node C^5^ had a k_ox_^O2^ of 160±10 M^-1^s^-1^ that was similar to that of Node C Thr300Glu (Table 1 and Fig. 6C), indicating that the additional four substitutions in this variant do not further enhance O_2_ reactivity beyond that of Thr300Glu despite the fact that individually introducing the “reverse” mutations into Node D produces a precipitous decline in activity in steady state assays (Fig.5). Node C^13^ had a k_ox_^O2^ of 3.7±0.3*10^3^ M^-1^s^-1^ (Table 1 and Fig. 6D), about 230 times greater than Node C, indicating that the collective introduction of 13 substitutions that decreased activity by more than 45% in the Node D➔C reverse mutants substantially accelerate the reaction with O_2_, though not quite to the level of Node D. However, Node C^21^, which includes all 21 substitutions that individually decreased activity by >25% in the Node D➔C “reverse” mutants, resulted in a variant with k_ox_^O2^ of 1.9±0.2*10^4^ M^-1^s^-1^ (Table 1 and Fig. 6D): this is a remarkable >1100-fold increase from that of Node C, making it as effective with O_2_ as other known oxidases from the FAO family. Notably, the k_ox_^O2^ value observed for Node C^21^ is nearly identical to the k_ox_^O2^ for Node C^55-oxidase^, indicating that the 21 substitutions capture the functionally most important amino acid replacements between Node C and Node D that gave rise to the potent oxidase activity in the NctB lineage. These results also suggest that most, if not all, of the 21 substitutions in Node C^21^ contribute to its gain in oxygen reactivity, indicating that control of oxygen reactivity is not confined to a few specific sites in this lineage of enzymes.

### Oxidase enhancing positions are distributed throughout the structure

We next sought to understand the structural basis for the enhanced oxidase activity between Node C and Node C^21^ by analyzing AlphaFold 3.0 models of the two proteins. Node C^21^ was selected for this analysis since it comprises the minimal number of amino acid changes needed to convert the Node C dehydrogenase into a potent oxidase. The reaction between flavin hydroquinone and O_2_ is thought to occur at the C4a/N5 region of the flavin isoalloxazine (5,13,17). However, the AlphaFold models show that the oxidase enhancing amino acid substitutions present in Node C^21^ are widely distributed throughout the protein structure with only two (Gly68GSer and Glu300Thr) having side chains less than 8 Å away from the flavin isoalloxazine (Fig. 7). This observation suggests that the majority of oxidase enhancing substitutions are not exerting their effects by modulating the local environment or reactivity of the isoalloxazine in the active site, but instead modulate O_2_ reactivity of the protein through long range effects like altering dynamics of the protein or O_2_ accessibility to the reaction center at the flavin C4a/N5.

**Fig. 7.**
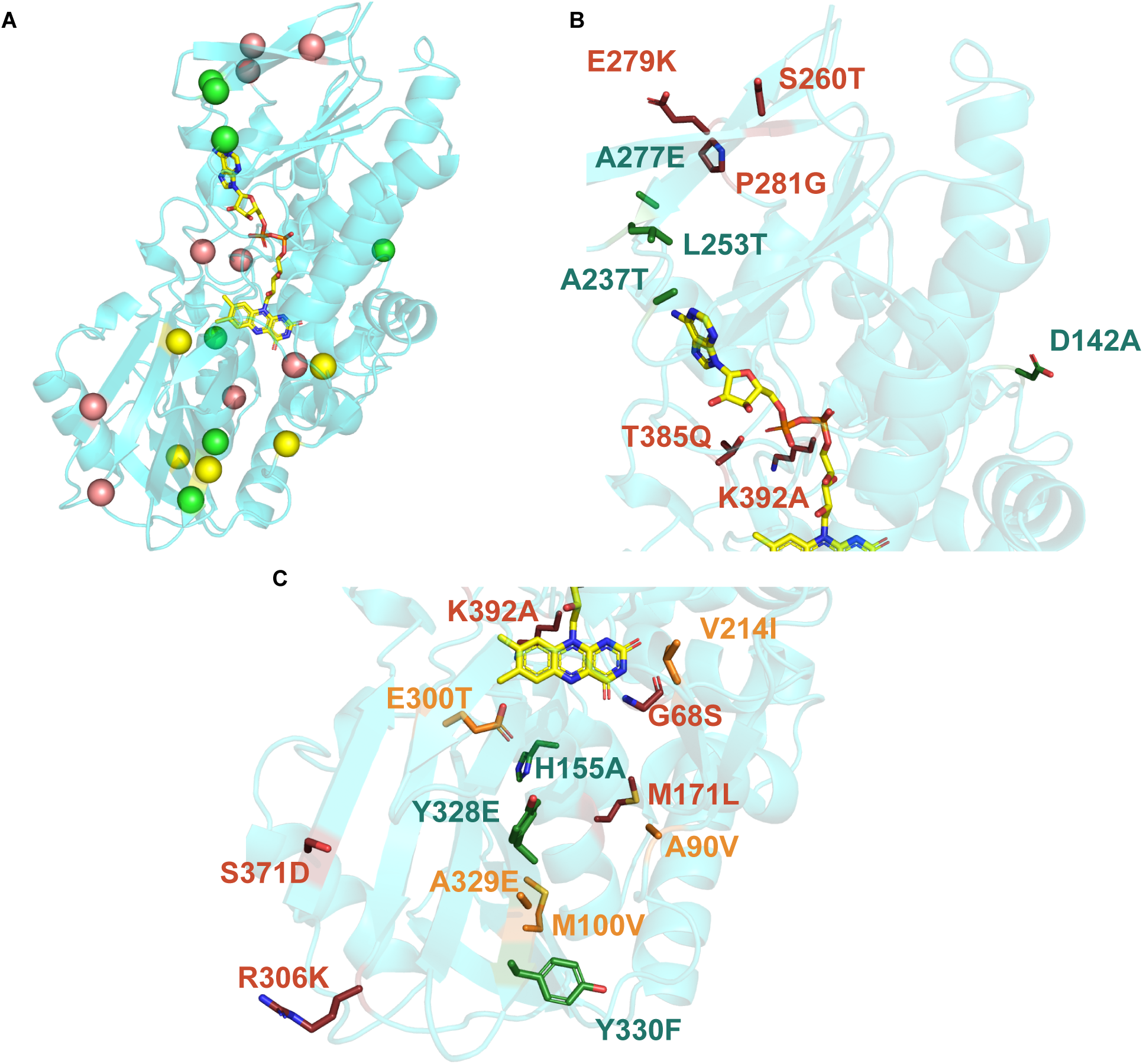
Residue positions that contribute to oxidase activity. **(A)** AlphaFold model showing a protomer of the predicted Node C^21^ dimer. The backbone is shown in cyan and the FAD group is shown in yellow sticks. Yellow, red and green spheres represent the C_α_ of residues that, when substituted, cause a >60%, 60-40%, and 40-25% decrease in oxidase activity, respectively, in the D➔C single mutants. **(B)** and **(C)**, Magnified view of Node C^21^ protomer focusing on the adenine and isoalloxazine ring-containing portions, respectively, of the Node C^21^ structure. Oxidase enhancing residues in Node C^21^ are shown as sticks. Labels for amino acid substitutions are written in the D➔C direction and are colored to be consistent with that shown in Figure 5.

### Molecular dynamics reveals major changes in mobility around the isoalloxazine

To further investigate the role of these oxidase-enhancing amino acid substitutions on protein dynamics, oxygen diffusion, and accessibility towards the flavin, we performed Gaussian accelerated molecular dynamics (GaMD) simulations on the Alphafold 3.0 models of Node C and Node C^21^ homodimers (34). GaMD adds a harmonic boost potential to the dihedral potential and overall potential of the system, accelerating the exploration of the free energy landscape. This results in a significant increase in the sampling of oxygen diffusion pathways. Three independent GaMD simulations were performed for Node C and Node C^21^, respectively, each lasting 500 ns. Oxygen diffusion was considered successful when the oxygen molecules reached within 6 Å of the C4A atom of the flavin isoalloxazine, which is the site of reaction with the flavin hydroquinone. A total of 59 successful independent diffusion trajectories were observed in Node C, and 58 successful independent diffusion trajectories in Node C^21^ across all replicates. Observed pathways were categorized into two groups: one approaching from the si-side of the flavin and the other from the re-side. The CAVER 3.0 plugin was used to visualize these diffusion pathways observed in the MD simulations (Fig. 8) (35). No significant differences were found in the routes by which oxygen arrives at the flavin isoalloxazine, suggesting that oxygen diffusion routes are not affected by the substitutions present in Node C^21^. On the si-side, several smaller pathways extend into the protein matrix and connect to the larger si-side solvent-exposed channel below the flavin isoalloxazine. The re-side pathways originate near the dimer interface and join an intermittently observed hydrophobic pocket located above the N1 atom of the flavin isoalloxazine (Fig. 8).

**Fig. 8.**
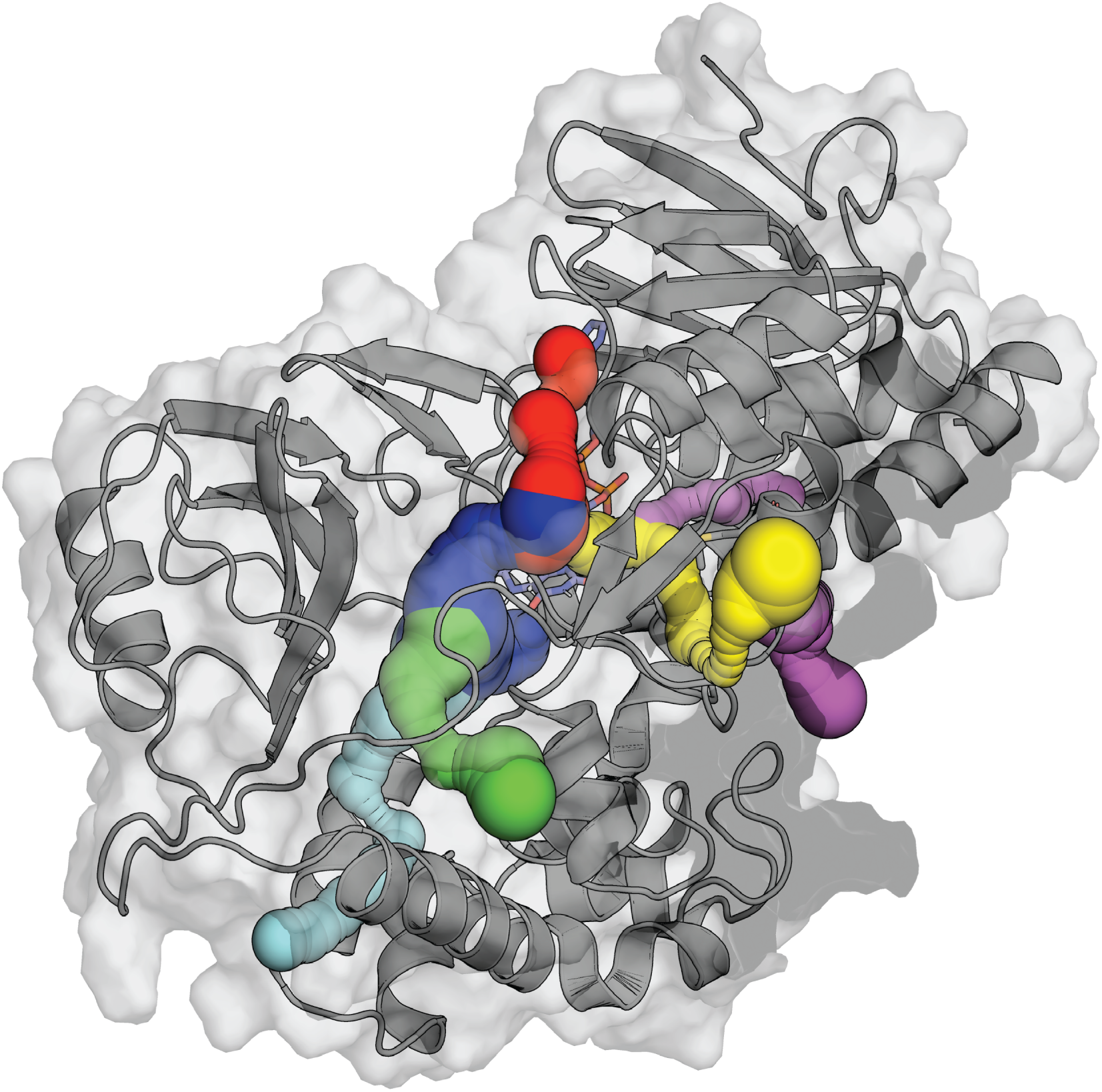
O_2_ diffusion pathways in Node C and Node C^21^. One chain of Node C^21^ highlighting the oxygen diffusion pathways towards the C4A atom on the flavin isoalloxazine as determined by GaMD simulations and visualized using CAVER3.0. Pathways colored cyan, red, blue and green show the most common si-side routes for oxygen transport. Pathways colored magenta and yellow show the most common re-side routes for oxygen transport. Pathways of oxygen diffusion were similar in the simulations of Node C.

Analysis of root mean square fluctuations (RMSF) of the backbone carbons revealed quantifiable differences between Node C and Node C^21^, indicating that global dynamics are altered by the presence of substitutions across the protein structure (Fig. 9). Node C exhibited an average RMSF of 24.43 Å compared to 13.53 Å for Node C^21^, indicating that the Node C^21^ oxidase is much less dynamic than the Node C dehydrogenase. This decrease in dynamics spanned the entire protein, with nearly every residue of the Node C^21^ homodimer having a lower RMSF than the corresponding position in Node C. Radius of gyration (Rg) measurements further quantified these differences, with Node C^21^ showing a more compact ensemble of structures (Mean Rg = 29.43 ± 0.15 Å) than Node C (Mean Rg = 29.84 ± 0.21 Å) (Fig. S8A). These data suggest that Node C^21^ has a less dynamic and more compact structure, which may contribute to its increased oxidase activity (Fig. S8B).

**Fig. 9.**
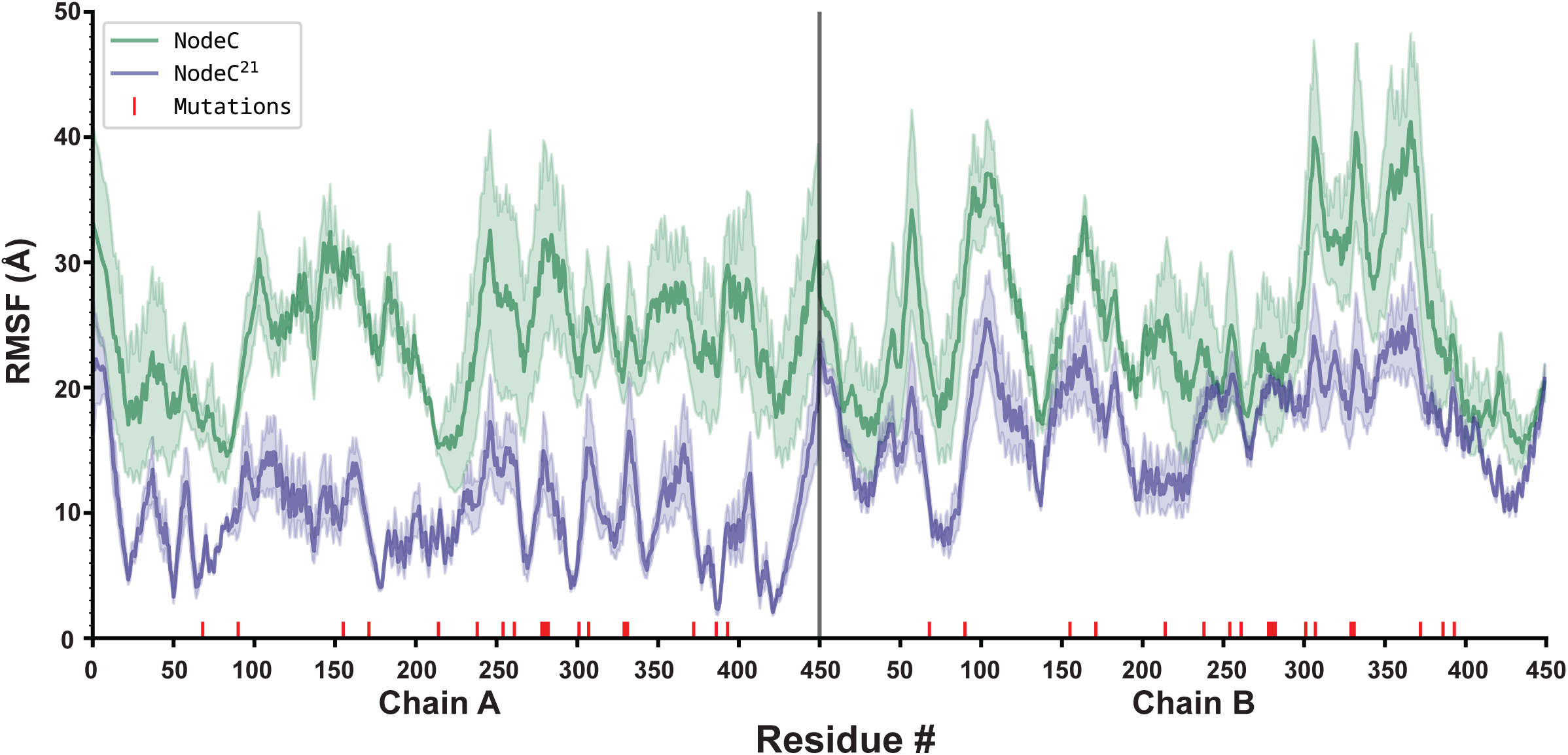
Flexibility of the protein backbone in Node C and Node C^21^. Root Mean Square Fluctuations (RMSF) of the backbone alpha carbons of Node C and Node C^21^ averaged over 3 independent runs. Node C (green) shows significantly higher RMSF compared to NodeC^21^ (purple) throughout, indicating a shift in global dynamics due to the 21 substitutions between these two proteins (marked in red).

Nearly all FAOs have a conserved lysine that forms a water-mediated hydrogen bond with the flavin N5, and this residue has been shown to be critical for accelerating the reaction with O_2_ in several oxidases of the FAO structural family (2, 3, 7, 17, 36–41). Lys340 is appropriately positioned to form a water-mediated hydrogen bond with the flavin N5 in the AlphaFold models of Node C, Node C^21^, and the crystal structure of NctB (Lys331), suggesting that Lys340 may accelerate the reaction of reduced flavin with O_2_ if Lys340 is correctly positioned. A detailed examination of the active site in the MD simulations revealed substantial differences in the position and mobility of Lys340 and surrounding residues between Node C and Node C^21^. The side chain of Lys340 resided significantly closer to the reactive center of the flavin isoalloxazine in Node C^21^, with an average distance of 8.11 ± 1.05 Å to the flavin N5 compared to 9.29 ± 1.33 Å in Node C (Fig. 10). This may, in part, be due to the more compact global ensemble of structures in Node C^21^, which appears to decrease the distance between Lys340 and flavin N5 in the MD simulations (Fig. S8C). The movement of Lys340 in Node C^21^ was also relatively static, with the ε-amino group of the side chain oriented towards the flavin N5 throughout the course of the simulation (Fig. 10 and Movie S1). In contrast, Lys340 displayed higher conformational flexibility in Node C, with the residue alternating between a position adjacent to the flavin N5 (the “in” conformation) and one where the side chain is flipped nearly 180 degrees away from the active site (the “out” conformation) to interact with Glu328 near the protein surface (Fig. 10 and Movie S2). Notably, Node C^21^ has a tyrosine at the position of this glutamate, and the Glu328Tyr substitution would be expected to disrupt the interaction with Lys340 in the “out” conformation observed in Node C. Residue-residue contact analysis using MDtraj indicated that this is indeed the case, with Lys340 forming a contact with Glu328 41% of the time in the simulations of Node C but only 7% of the time with Tyr328 in the simulations of Node C^21^ (Table S2). Residue-residue contact analysis indicated that the Thr300Glu substitution also likely contributes to the reduced mobility of Lys340 in Node C^21^ through polar interactions between the lysine and glutamate side chains (Movie S2), as Lys340 forms contacts with Glu300 96% of the time in Node C^21^ but contacts Thr300 only 43% of the time in the MD simulations of Node C (Table S2). Moreover, hydrogen bond analysis using cpptraj showed an almost 37-fold increase in hydrogen bond persistence between Lys340 and Glu300 throughout the simulation in Node C^21^ (Fig. S9). The Thr300 residue in Node C is unable to consistently form this interaction with Lys340, likely resulting in increased time that Lys340 spends in the “out” conformation.

**Fig. 10.**
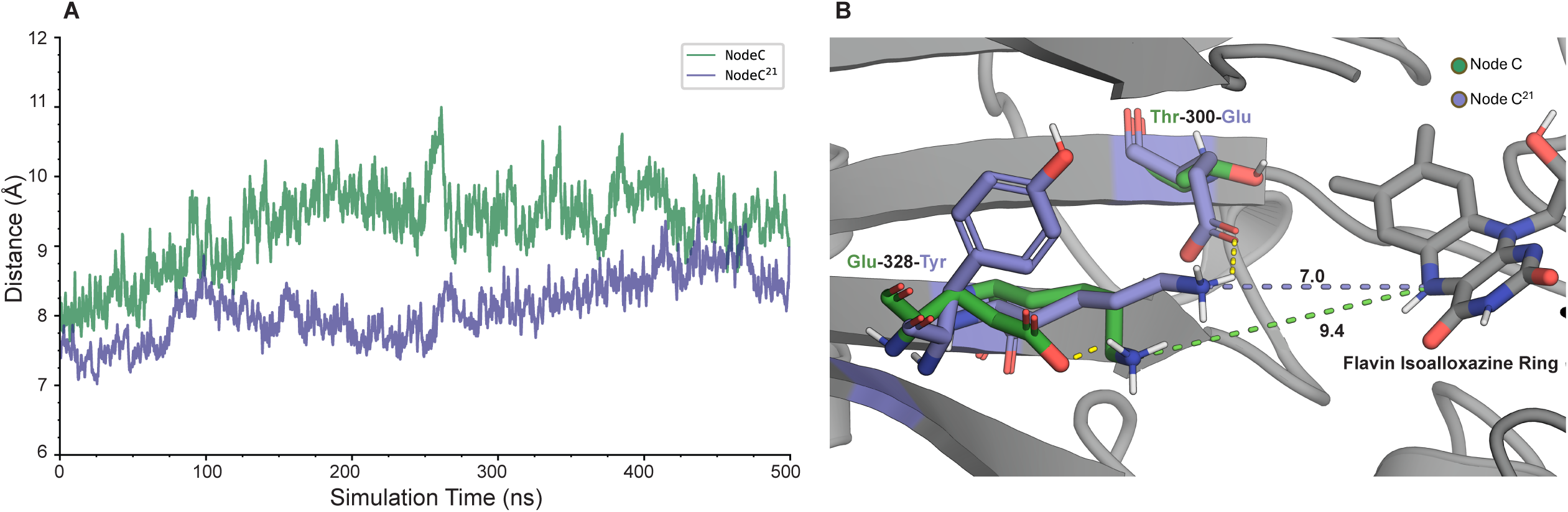
Lysine positioning in Node C and Node C^21^. (A) Distance between the side-chain ε-nitrogen of Lysine 340 and the N5 nitrogen on the flavin isoalloxazine over a 500 ns simulation, showing consistently shorter distances in Node C^21^ (purple) compared to Node C (green). (B) Structural comparison of the key residues in Node C (green) and NodeC^21^ (purple) at a representative snapshot from the MD simulations, highlighting the mutations (Thr300Glu and Glu328Tyr between Node C and Node C^21^) and their effect on polar contacts (yellow dotted lines) that influence the conformational positioning of Lys340 relative to the flavin isoalloxazine. The distances illustrate their effect on the position of Lys340.

## Discussion

Here, we used a historical approach to dissect the evolution of oxidase activity in a set of nicotine/6OHN degrading FAOs and systematically identified the amino acid substitutions responsible for this gain in O_2_ reactivity. The tandem genomic association of cytochrome c genes with *nicA2* and *nctX* genes suggests that the ancient ancestor that gave rise to extant NicA2, NctX, and NctB proteins was a dehydrogenase (Fig. 1), and this is supported by the low reactivity of Node A with O_2_ and its rapid kinetics of oxidation using CycN as oxidant. Our results indicate that the rapid reactivity with O_2_ of extant NctB enzymes evolved from within this larger group of dehydrogenases via a subset of the amino acid changes between Node C and Node D. Without functional characterization of other, more distantly related bacterial FAO homologs, it is difficult to predict the O_2_ reactivity of ancestors that preceded Node A. However, the widespread absence of tandem genomic cytochrome c genes among bacterial FAO homologs suggests that the ancient-most ancestral enzymes that existed prior to the evolution of Node A were oxidases. This is consistent with phylogenetic analyses of the FAO superfamily showing that characterized members of every other lineage of related enzymes are oxidases (42). This apparent ancestral oxidase activity was lost when the Node A enzyme became a dehydrogenase, only to be regained in the NctB lineage in the form of an evolutionary reversal between Node C and Node D.

Our results highlight the value of ASR as an efficient approach to elucidate the structural and molecular bases underlying changes in enzymatic function. Specifically, ASR enabled us to drastically reduce the number of mutations to investigate, as only 77 amino acids were replaced over the historical episode during which oxidase activity evolved between Nodes C and D (Fig. 1A). In constrast, if we had only directly compared differences between the modern-day NicA2 dehydrogenase and NctB oxidase, we would have had 242 sites to evaluate. Our systematic biochemical analysis of substitutions between Nodes C and D identified a minimal set of 21 amino acid replacements during the divergence of these two proteins that gave rise to the potent O_2_ reactivity observed in the Node D oxidase. Most of these substitutions are distant from the flavin isoalloxazine, with several of them even having their side chains oriented towards the solvent on the surface of the protein in AlphaFold models.

These 21 substitutions that enable rapid reaction with O_2_ do not appear to significantly alter O_2_ diffusion through the protein matrix, but instead result in a large scale, global decline in the conformational flexibility of the enzyme (Fig. 9). This decrease in global dynamics is associated with a significant restriction in the movement of the active site, in particular that of Lys340 near the flavin N5 (Fig. 10). Mechanistic and structural work on several oxidases of the FAO superfamily has demonstrated that a conserved lysine residue proximal to the flavin N5 is critical for the ability of these enzymes to react rapidly with O_2_ (2, 3, 7, 15, 34–39, 41). Oxidation of flavin hydroquinone by O_2_ requires an initial single electron transfer step to form flavin semiquinone and negatively-charged superoxide (O_2_ ^−^) prior to spin inversion and a second single electron transfer to yield oxidized flavin and hydrogen peroxide. The initial single electron transfer to form O_2_ ^−^ is the slow step in this process, which may be facilitated by the positive charge afforded by the conserved lysine to stabilize the transition state for forming O_2_ ^−^, ultimately accelerating the reaction with O_2_ (7–9). Lys340 is appropriately positioned in the structures of Node C and Node C^21^ to fulfill this role, but is conformationally dynamic in our simulations of Node C, with the side chain of Lys340 alternating between the “in” conformation oriented towards the flavin and the “out” conformation where it is flipped away from the flavin to interact with Glu328 near the surface of Node C (Fig. 10). In the “out” conformation, it would be incapable of facilitating the reaction with O_2_, which is consistent with the very low k_ox_^O2^ value of the Node C enzyme. In contrast, Lys340 is less mobile in the simulations of Node C^21^ and is oriented with its ε-amino group towards the flavin throughout the course of the simulation (Fig. 10). The conformation of Lys340 in Node C^21^, which is also closer to the flavin on average than in Node C, is presumably more optimal to promote the reaction with O_2_, resulting in the dramatic acceleration in kinetics of oxidation in Node C^21^ compared with that of Node C.

The Thr300Glu substitution between Node C and Node C^21^ likely plays a key role in the conformational differences observed in Lys340, as the side chain of Glu300 in Node C^21^ would form a favorable charge-charge interaction with the Lys340 side chain that would be expected to stabilize Lys340 in the “in” conformation oriented towards the flavin. However, the Thr300Glu substitution is clearly not the only switch responsible for the change in oxygen reactivity, as the single Thr300Glu substitution into Node C increases k_ox_^O2^ by only 9-fold compared to the nearly 1100-fold improvement in k_ox_^O2^ upon introducing all 21 substitutions in Node C^21^ (Fig. 6C&D and Table 1). Similarly, the Glu328Tyr substitution also appears to contribute to the change in dynamics of Lys340 since the side chain of Lys340 interacts with the side chain of Glu328 in the simulations of Node C when the lysine is in the “out” conformation. Mutating Glu328 to tyrosine, as observed in Node C^21^, would disrupt this interaction, biasing Lys340 to spend more time in the “in” conformation adjacent to the flavin N5 necessary for promoting the reaction with O. While the side chains in the Thr300Glu and Glu328Tyr substitutions interact directly with Lys340 and bracket the two conformational extremes of Lys340, there are global changes in the structure and dynamics of the two proteins, with Node C^21^ being much less conformationally dynamic and more compact overall than Node C (Fig. 9 and Fig. S8). The substitutions at the other 19 sites changed between Node C and Node C^21^ presumably contribute to this global compaction and decreased conformational flexibility of the enzyme. These global changes would likely also restrict the flexibility and positioning of the active site and Lys340, contributing to the decreased average distance observed between the side chain of Lys340 and the flavin C4a-N5 in Node C^21^ (Fig. 10 and Fig. S8). The net effect of these global changes would therefore be to further optimize the positioning of Lys340 to promote the reaction with O_2_, leading to the dramatic acceleration in kinetics of oxidation observed in Node C^21^ relative to Node C. Further investigation will be needed to pinpoint the precise roles of these distal substitutions in modulating the global dynamics and active site fluctuations.

Flavoenzymes have the remarkable ability to tune the reactivity of their flavin cofactors with O_2_, with some oxidases having k_ox_^O2^ values near 10^6^ M^-1^s^-1^ and some dehydrogenases with k_ox_^O2^ values approaching 0 M^-1^s^-1^ (6, 9, 44). Decades of mechanistic work have focused on elucidating the structural and mechanistic features that enable flavoprotein oxidases to react efficiently with O_2_, with most studies understandably focusing on the local environment of the flavin cofactor (e.g., Glucose oxidases, Berberine bridge enzyme V.S. Pollen allergen Phl p 4) (45, 46). In these studies, a positive charge and nonpolar site proximal to the O_2_ reaction center are recurring features shown to be important in promoting the reaction of flavin hydroquinone with O_2_ in flavoprotein oxidases (6–9). However, it is considerably less clear why many flavoprotein dehydrogenases react so slowly with O_2_, particularly for dehydrogenases homologous to and sharing a highly conserved active site with oxidases (e.g., flavocytochrome b2 and glycolate oxidase) (4, 9, 47). O_2_ must access the flavin C4a/N5, and there is evidence of defined O_2_ diffusion pathways and specific gatekeeper residues that control O_2_ access in specific flavoenzymes (5, 10, 48–54). Our findings here show that the switch between dehydrogenase and oxidase function in the NctB lineage is brought about by a contellation of amino acid changes that fine-tune the positioning and conformational space and dynamics of a lysine residue adjacent to the flavin. In the dehydrogenase, this lysine residue is afforded considerable conformational freedom, allowing it to adopt conformations that are poorly suited for catalyzing the reaction with O_2_, ultimately suppressing this reaction in the dehydrogenase. In the oxidase, the lysine is restricted to the flavin-oriented conformation, placing its side chain in close proximity to where the reaction with O_2_ occurs, thus accelerating this reaction. Most of the amino acid replacement sites that contribute to the gain in oxidase function are distant from the flavin and do not directly interact with this lysine residue, but instead appear to exert their effects by modulating the global flexibility and structural space of the enzyme. These results therefore demonstrate that multiple amino acid substitutions far from the flavin isoalloxazine can have a profound impact on how efficiently the cofactor reacts with O_2_, a result which may apply to other enzyme families.

## Materials and Methods

### Phylogenetic tree construction and ancestral sequence resurrection

The FAO dataset used was constructed as follows. The previously experimentally characterized NctB sequence from *Shinella* sp. HZN7 was used as an query sequence in BLASTp search in GenBank non-redundant protein sequences (nr), carried out in May 2022. The 500 top hits were downloaded as aligned sequences from the BLAST search result to eliminate the N-terminal signal peptides from the dataset. This was done because the signal peptides are not part of the catalytic domain of the enzymes and not all of the sequences contained signal peptides. Accordingly, the signal peptides were removed to prevent them from biasing the subsequent phylogenetic analysis. An alignment of these sequences was achieved using MAFFT version 7 with the FFT-NS-2 strategy (26, 55). The sequence alignment was then used as input in IQ-Tree version 2.1.2 for phylogenetic tree construction and ancestral sequence prediction via the CIPRES science gateway, with IQ-Tree automatically selecting the best substitution model (LG+F+I+G4) (27, 56). Bootstrapping was carried out with 1000 ultrafast bootstraps within IQ-Tree. The output ancestral sequence predictions for the selected nodes were compared and reviewed with the input sequence alignment using Aliview to remove insertions in the ancestral reconstructions resulting from outlier sequences using parsimony criteria (57). Ancestral reconstruction constructs began at the start of the FAO structural domain with a DYD(A/V)IV motif at the N-terminal end. An unstructured peptide consisting of MTKAGHSASGEA, was included on the N-terminus of all ancestral reconstructions to promote stability of the ancestral reconstructions. This short peptide was derived from the predicted disordered N-terminal region preceding the core structural domain of the Node C ancestral reconstruction.

### Protein expression and purification

The genes for the enzymes used in this study were synthesized by Twist Biosciences and cloned into the NdeI and XhoI sites of pET29b to generate constructs with a C-terminal His tag. Subsequently, the vector was introduced into *E. coli* BL21 (DE3) cells and cultivated in expression media consisting of 12 g/L tryptone, 24 g/L yeast extract, 40 ml/L glycerol, 0.072M K_2_HPO_4_, and 0.017M KH_2_PO_4_ (PEM). Cultures were incubated at 37 °C with shaking until an OD_600_ of 0.8. For FAOs the temperature was lowered to 20 °C, and cultures were induced with 100 μM IPTG and left to grow overnight. For CycN the temperature was lowered to 30 °C, and cultures were induced with 10 μM IPTG and left to grow overnight. Pellets were harvested and resuspended in lysis buffer (300 mM NaCl, 50 mM NaH_2_PO_4_, 20 mM imidazole, 10% v/v glycerol, pH 8). The resuspended cells were lysed by sonication after EDTA-free protease inhibitor cocktail (Abcam) and benzonse nuclease (Sigma) were added. The lysate was cleared by centrifugation at 24000 x g for 25 minutes. Supernatant was collected and loaded onto a nickel affinity column (resin from GBiosciences) that had been pre-equilibrated with lysis buffer. The column then was washed with 10 column volumes (CV) of lysis buffer and proteins were eluted with 2 CV of elution buffer (300 mM NaCl, 50 mM NaH_2_PO_4_, 250 mM imidazole, 10% v/v glycerol, pH 8). Proteins were concentrated and then subjected to an additional purification by passing through a HiLoad 16/100 Superdex 200 pg (75pg for CycN) column at 4° C. This purification process was carried out using buffer consisting of 40 mM Hepes, 100 mM NaCl, and 10% v/v glycerol, at pH 7.5 (stopped-flow buffer). The purified FAOs were concentrated and flash frozen for storage at -80° C. Purified CycN was stored at 4° C.

### Transient kinetic assays

All transient kinetics experiments were done in stopped-flow buffer at 4° C using a TgK Scientific SF-61DX2 KinetAsyst stopped-flow instrument. All protein solutions were made anaerobic in glass tonometers by repeated cycles with vacuum and anaerobic argon (58). When needed, the FAD group in FAOs was reduced in the anaerobic tonometer by titrating with 1mg/mL dithionite (for Node A) in stopped-flow buffer or 20 mM anaerobic 6OHN (for all enzymes other than Node A) in a gas tight hamilton syringe. Anaerobic 6OHN was prepared by sparging for at least 10 min with anaerobic argon. The dithionite or anaerobic 6OHN was slowly added to the point where the FAD group was fully reduced to the hydroquinone state, with the titration process monitored spectrophotometrically using a Cary 50 Bio UV–Visible Spectrophotometer and Cary WinUV software.

For experiments monitoring the reaction with O_2_, the FAO containing FAD hydroquinone in a tonometer was loaded on the stopped-flow instrument and mixed with buffer containing various oxygen concentrations (prepared by sparging buffer with various N_2_/O_2_ ratios made using a gas blender). For experiments monitoring the reaction with CycN, another tonometer with various concentratrations of anaerobic CycN was loaded on the stopped flow instrument. These reactions were monitored using the instrument’s multiwavelength charge-coupled device detector. For experiments monitoring the reaction with nicotine or 6OHN, oxidized anaerobic FAOs was mixed with various concentrations of nicotine or 6OHN (made anaerobic by sparging with argon), and the reaction was monitored using the instrument’s single-wavelength photomultiplier tube detector at 450 nm.

Stopped-flow data were analyzed using Kaleidagraph. Kinetic traces at 450 nm for the reaction with oxygen for all FAOs were fit to a single exponential function (Equation 1) to determine the observed rate constant (k_obs_) for each oxygen concentration. Traces for the reaction of Node B reacting with nicotine were also fit to Equation 1. Kinetic traces for all other experiments were fit to a sum of two exponentials (Equation 2) or a sum of three exponentials for reaction with CycN (Equation 3) to determine k_obs_ values for the various kinetic phases. The kinetic amplitude for each phase is represented by ΔA, the apparent first-order rate constant is denoted as k_obs_, and A_∞_ refers to the absorbance at the completion of the reaction.

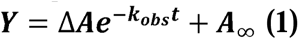

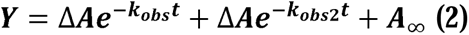

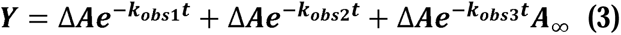

Plots of k_obs_ against substrate concentrations for reaction with oxygen or CycN were linearly fitted, with the slope providing the second-order rate constant between the reaction of reduced enzyme and oxidant (k_ox_). Plots of k_obs_ against nicotine and 6OHN displayed a hyperbolic dependence and was fit to Equation 4 to determine k_red_, the rate constant for flavin reduction and K_d_ for substrate binding to oxidized FAO.

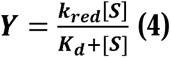

### Site-directed mutagenesis

Single mutants of Node C ASR were prepared via site-directed mutagenesis using PCR and mutagenic primers (59). The introductions of mutations were verified by Sanger sequencing and the mutants were purified using the same protocol described above.

### Steady state kinetics for oxidase activity screening

All experiments were completed in stopped flow buffer at 4°C using a Hansatech Oxygraph+ oxygen probe system to monitor the rate of oxygen consumption. Stopped flow buffer containing 200 µM 6OHN was placed in the chamber of the system. FAO variant was then injected into the chamber using a Hamilton syringe to make the final concentration 200 nM. The rates of oxygen consumption were recorded and the consumption rate of Node C was set as 100% oxygen reactivity. Percentage of oxygen reactivity of other mutants and Node C were calculated as equation shows.

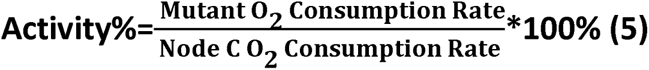

### Modeling structures using AlphaFold3 monomer and multimer functions

Node B and Node C structures were modeled using AlphaFold 3.0 in the Alphafold server using the casp14/full_dbs settings with both monomer and multimer functions, with two copies of FAD included as ligand in the generation of the AlphaFold models (60).

### Molecular dynamics simulations and analysis of O_2_ diffusion

The PDB files of the AlphaFold 3.0 structures of Node C and Node C^21^ were processed and prepared using Schrödinger Maestro (34). Missing hydrogens were added, and protonation states of titratable residues were set at pH 7.4 using the Protein Preparation Tool. Two hundred oxygen molecules were inserted randomly, replacing bulk water molecules using Packmol (61). Atomic point charges for the non-standard flavin adenine dinucleotide (FAD) molecule were computed using the Antechamber tool in the AMBER22 software suite (62). The ff19SB protein forcefield was used to parametrize the standard residues, while the General Amber Force Field 2 (Gaff2) was used for non-standard residues. The tleap module in AMBER22 was used to solvate the protein with the OPC water model.

Molecular dynamics simulations were performed using the AMBER22 software suite. Energy minimization was conducted using the steepest-descent algorithm, first on the heavy atoms and then on the entire protein. The system was heated from 10 K to 300 K over 10 ns, followed by equilibration in multiple relaxation steps for a total of 20 ns, transitioning from the NVT ensemble to the NPT ensemble. To ensure sufficient exploration of oxygen diffusion pathways, enhanced sampling via Gaussian accelerated molecular dynamics (GaMD) was employed (34). Three independent GaMD simulations were performed for Node C and Node C^21^, each lasting 500 ns.

Post-processing of the simulation output for root mean square fluctuation (RMSF), radius of gyration (Rg), distance, and hydrogen bond analyses was performed using the cpptraj module in AMBER22. Oxygen diffusion pathways were validated using the CAVER 3.0 plugin (35). Residue-residue contact map analysis was conducted using MDtraj (63). Graphs and further processing of analyzed data were generated using Python 3.11. Figures were made using Pymol 3.0 and movies of simulation trajectories were recorded using ChimeraX (64).

## Data Availability

All data are contained within the article.

## Supporting Information

This article contains supporting information.

## Funding and Additional Information

This research was supported by National Science Foundation grant 2236541 (to F. S.) and the University of California, Davis, Department of Chemical Engineering start-up funds (to S. A.). P.R.B acknowledges support from NSF ACCESS BIO240267.

## Acknowledgments

P.R.B. thanks Michael Toney for giving helpful suggestions and feedback. All simulations were done using the San Diego Supercomputing Cluster (SDSC) and HPC2 core facility at University of California, Davis.

## Author Contributions

F.S. and T.J.B. conceptualization; Z.Z., P.B., S.A., T.J.B. and F.S. methodology; Z.Z., P.B., and A.G. investigation; Z.Z., P.B. and F.S. visualization; F.S. and S.A. supervision; Z.Z., P.B. and F.S. writing–original draft; Z.Z., P.B., S.A., T.J.B. and F.S. writing–review & editing.

## Abbreviations–The abbreviations used are

FAO: flavoprotein amine oxidase;
FADH_2_: reduced FAD hydroquinone;
O_2_: dioxygen;
k_ox_^O2^: second order rate constant for reaction with O_2_;
NicA2: nicotine oxidoreductase;
CycN: cytochrome c from *Pseudomonas putida* S16;
NctB: (S)-6-hydroxynicotine oxidase from *Shinella* sp. HZN7;
ASR: ancestral sequence reconstruction;
NctX: NctB-like enzymes from the Sphingomonadaceae family of bacteria;
6OHN: (S)-6-hydroxynicotine;
k_red_: rate constant for flavin reduction;
k_obs_: observed rate constant;
CycN_ox_: oxidized CycN;
k_ox_^CycN^: second order rate constant for reaction with CycN;
C➔D: a mutation in the Node C to Node D direction;
D➔C: a mutation in the Node D to Node C direction;
Node C^55-oxidase^: variant of Node C containing the 55 substitutions that uniquely changed between Node C and Node D;
Node C^22-dehydrogenase^: variant of Node C containing the 22 substitutions that did not uniquely change between Node C and Node D;
Node C^5^: variant of Node C containing the following five substitutions: V90A, T300E, V100M, E329A and I214V;
Node C^13^: variant of Node C containing the following thirteen substitutions: V90A, T300E, V100M, E329A, I214V, G281P, Q385T, S68G, A392K, T260S, K279E, L171M, K306R and D371S;
Node C^21^: variant of Node C containing the following 21 substitutions: V90A, T300E, V100M, E329A, I214V, G281P, Q385T, S68G, A392K, T260S, K279E, L171M, K306R, D371S, A142D, E328E, E277A, T253L, A155A, T237T and F330Y;
GaMD: Gaussian accelerated molecular dynamics;
RMSF: root mean square fluctuation;
R_g_: radius of gyration.

## Notes

### Competing Interest Statement

The authors have declared no competing interest.

