## Supplementary information for "Ancestral evolution of oxidase activity in a class of (S)-nicotine and (S)-6-hydroxynicotine degrading flavoenzymes"

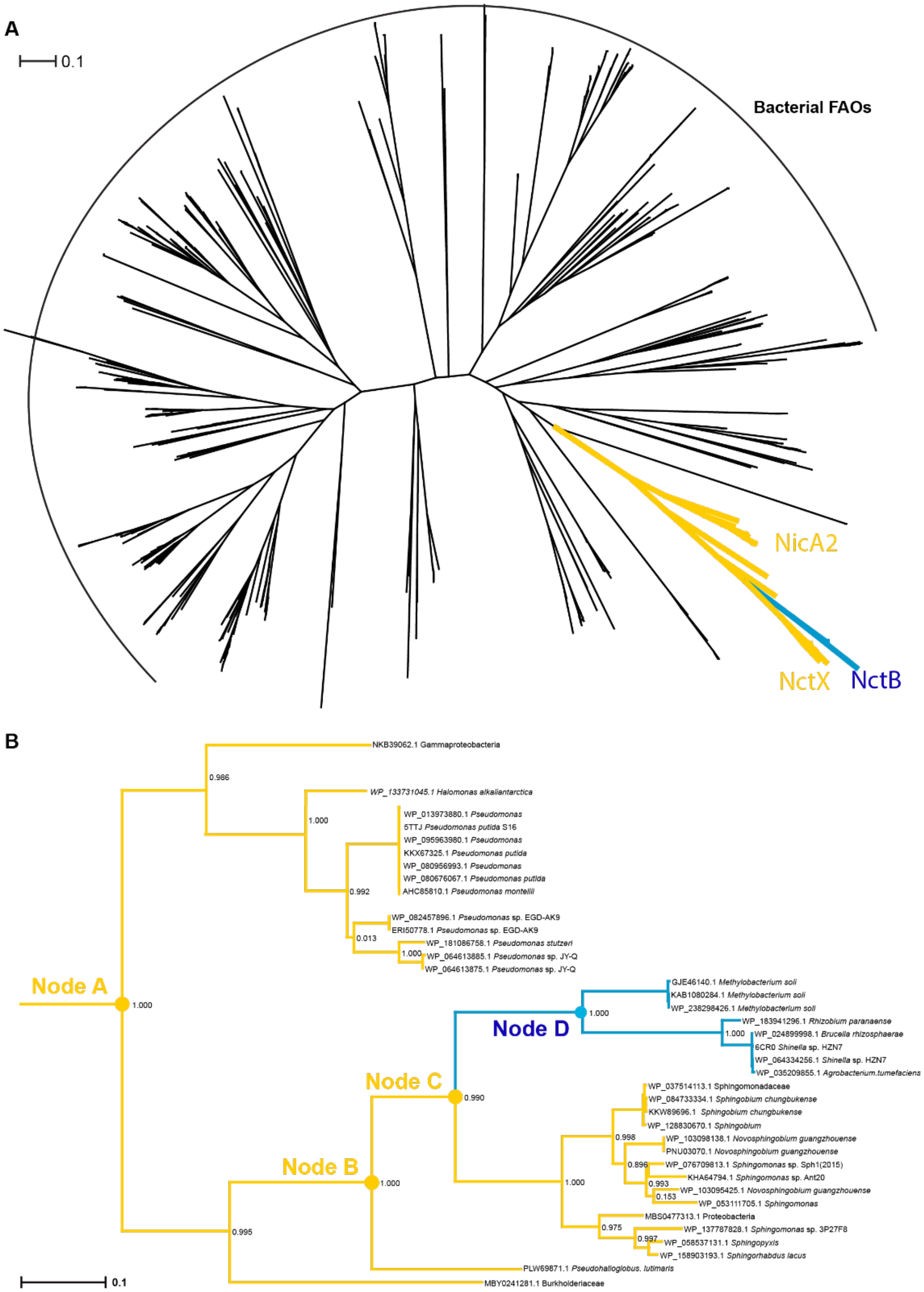


**Fig. S1.** Phylogeny of bacterial FAOs. (A) Radial phylogram of 500 bacterial FAO sequences obtained from a BLAST search using NctB as a query. Sequences were aligned using MAFFT and the tree generated using IQTree (1,2). The clade of interest in this study is colored in yellow (NicA2 and NctX) and blue (NctB). (B) Phylogram showing the detailed clade of interest with species names and GenBank accession numbers. Branches are colored yellow if the descendant FAO sequences are found in an operon with a cytochrome c and are colored blue if the descendant FAO sequences lack an adjacent cytochrome c.

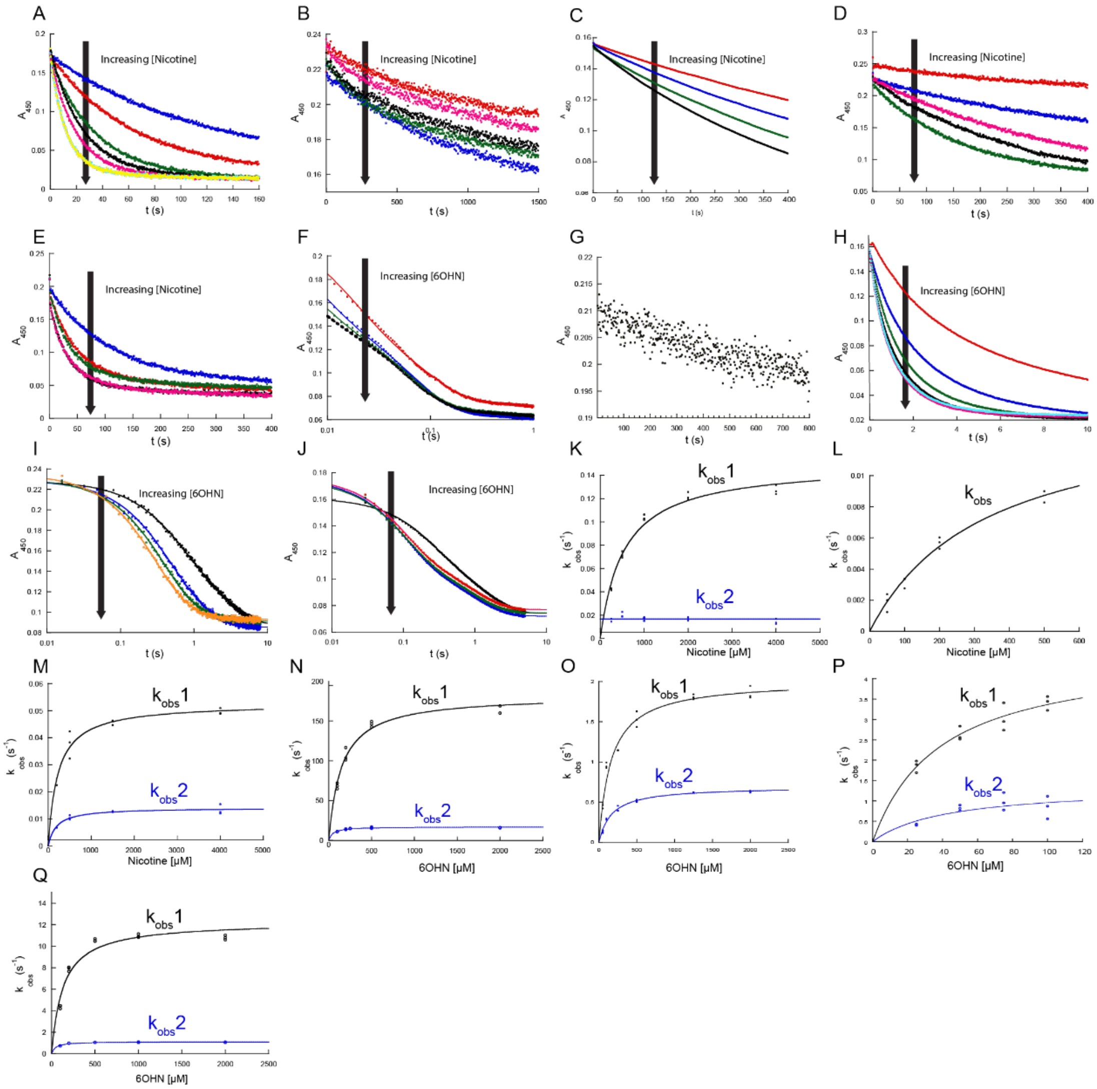


**Fig. S2.** Kinetic traces and parameters of flavin reduction by nicotine or 6OHN for NctB and ancestral enzymes. (A), (B), (C), (D) and (E), Absorbance trace overlay at 450 nm for the reduction of NctB, Node A, Node B, Node C, and Node D, respectively, by various concentrations of nicotine. Node B traces were fitted to equation 1. NctB, Node C traces can be fit to equation 2. Node A traces are too slow for accurate fitting. (F), (G), (H) and (I), and (J) Absorbance trace overlay at 450 nm for the reduction of NctB, Node A, Node B, Node C and Node D, respectively, by various concentrations of 6OHN. Only one concentration of 6OHN was shown for Node A given the slow reaction with 6OHN. Note that except for Node A, others have logarithmic time scales and can be fit to equation 2. (K), (L) and (M), $k_{obs}$ values for the first (black) and second (blue) phase plotted against the concentration of nicotine for NctB, Node C and Node D, respectively. Observed rate constants were plotted against nicotine concentration and the plot was fitted to Equation 4. The $k_{obs}$ value for the second phase was invariant with nicotine concentration for NctB. (N), (O), (P) and (Q), $k_{obs}$ values for the first (black) and second (blue) phase plotted against the concentrations of 6OHN for NctB, Node B, Node C, and Node D respectively. Both phases show hyperbolic dependence and were fitted to Equation 4. The $k_{red}$ and $K_d$ values associated with these data sets can be found in Table S1.

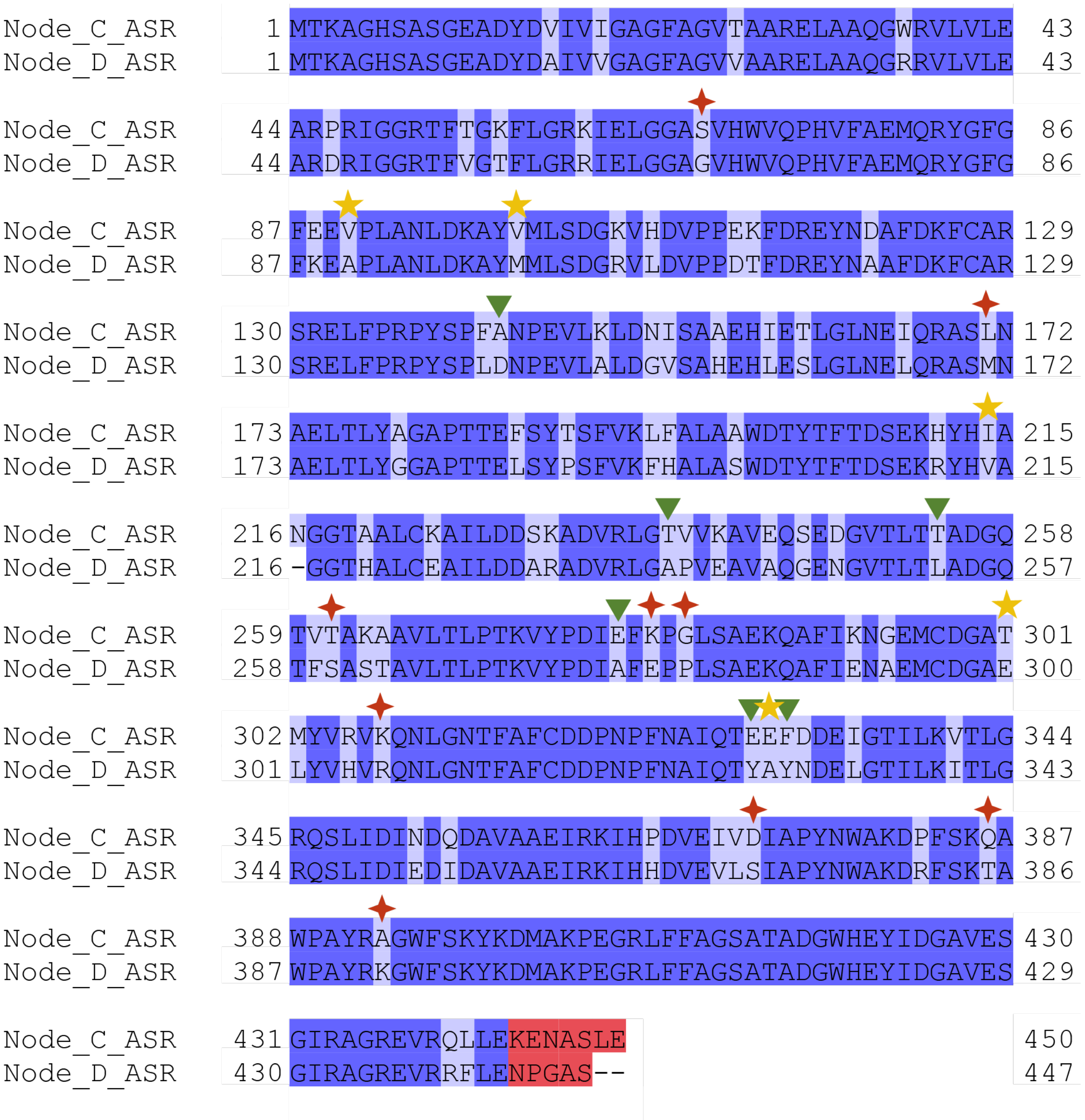


**Fig. S3.** Alignment of Node C and Node D protein sequences. Yellow Stars, red four-pointed stars, green triangles denote amino acid locations causing a >60%, 40-60%, and 40-25% decrease in oxidase activity, respectively. Sequences are colored to show the identity of residues. The 5-7 amino acids at the C-terminus (colored red) are predicted to be disordered and not part of the protein's core structural domain and were therefore not investigated for their impact on oxidase activity.

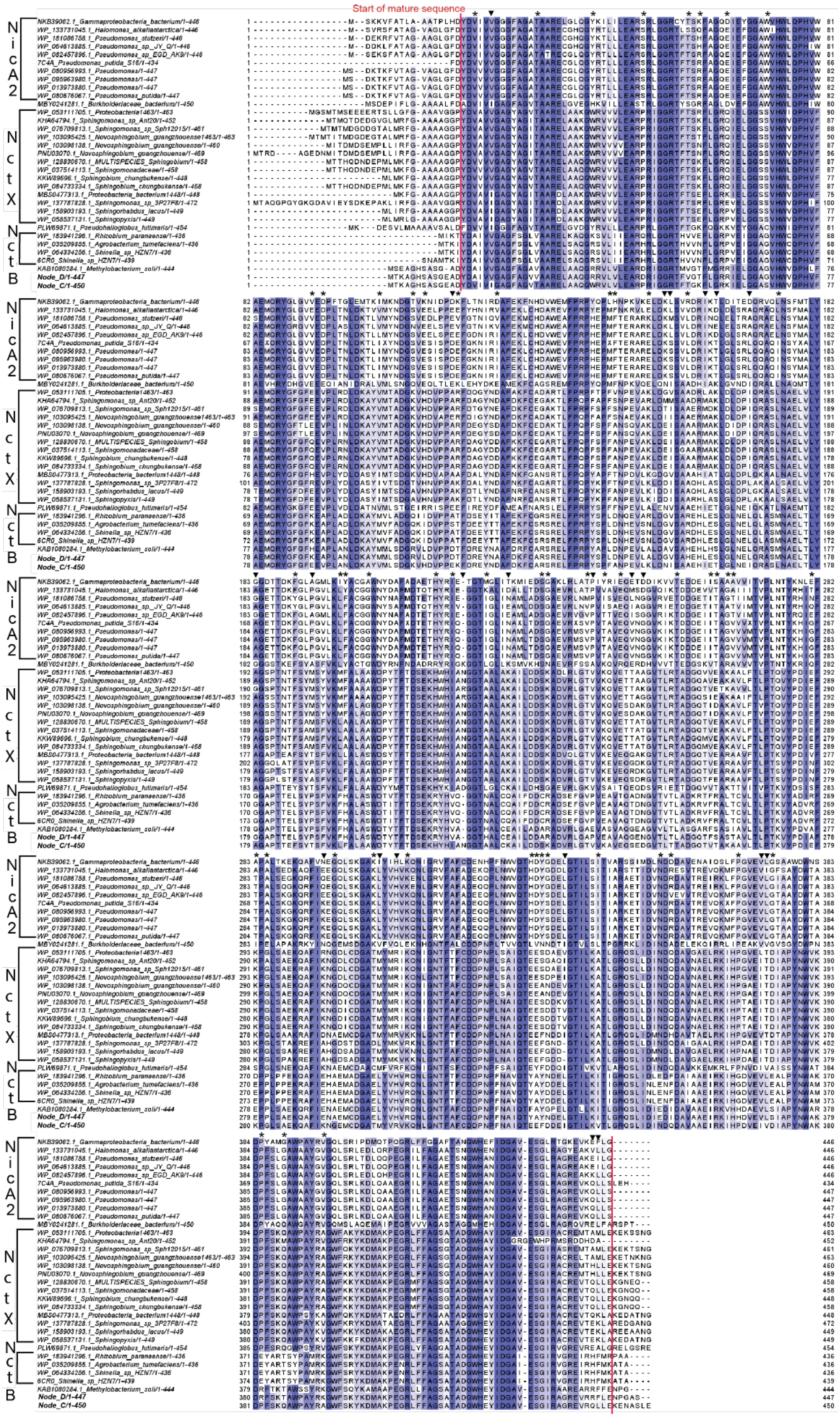
Start of mature sequence
NicA2
NctX
NctB

**Fig. S4**. Alignment of sequences in clade of interest used for screening residues included in Node $C^{55\text{-oxidase}}$ and Node $C^{22\text{-dehydrogenase}}$. Asterisks denote amino acid positions replaced in Node $C^{55\text{-oxidase}}$. Triangles denote amino acid positions replaced in Node $C^{22\text{-dehydrogenase}}$. If an amino acid residue present in Node D at a given position was also found in some of the NicA2 or NctX dehydrogenases, it was considered a non-unique site and was included in Node $C^{22\text{-dehydrogenase}}$. For example, at site 19 (Node D numbering), Node C contains isoleucine whereas Node D contains valine. However, valine is also present at this position in several NicA2 and NctX dehydrogenase sequences and we therefore hypothesized that the isoleucine to valine substitution at this position was not important for evolution of oxidase activity between Node C and Node D. Sites where the Node D residue was not found in any of the NicA2 or NctX sequences were considered unique sites and were included in Node $C^{55\text{-oxidase}}$. For example, at site 16 (Node D numbering), Node C contains valine and Node D contains alanine; alanine is not found at this site within any of the NicA2 or NctX dehydrogenase sequences, making it a unique change in the Node C to Node D lineage and was therefore included in Node $C^{55\text{-oxidase}}$. Residues are colored for their degree of conservation using the BLOSUM62 score. The figure was prepared using Jalview.

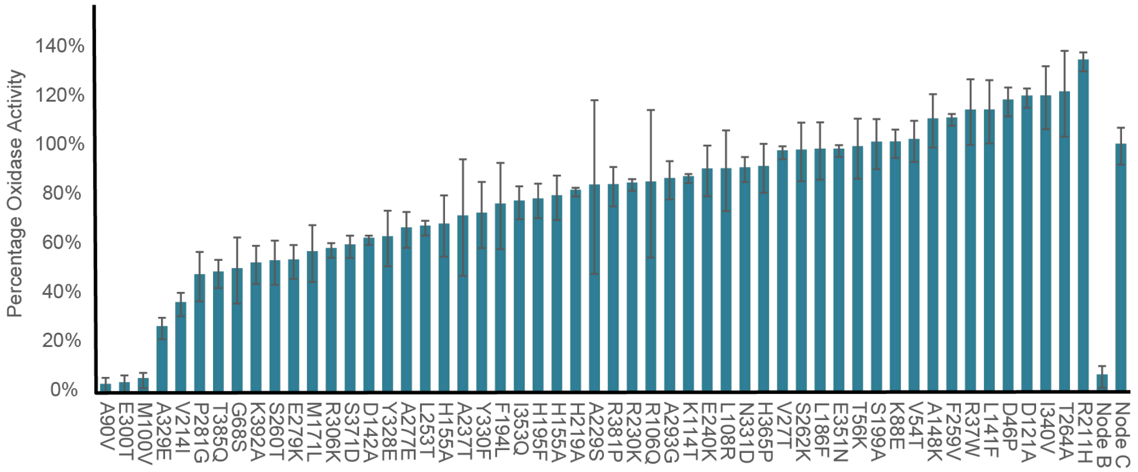


**Fig. S5.** Relative oxidase activity of all D→C single mutants tested. Numbers represent the residue position in Node D and replacement information indicates the amino acid in Node D that is replaced with the residue present in Node C for each D→C variant. Error bars represent standard deviations.

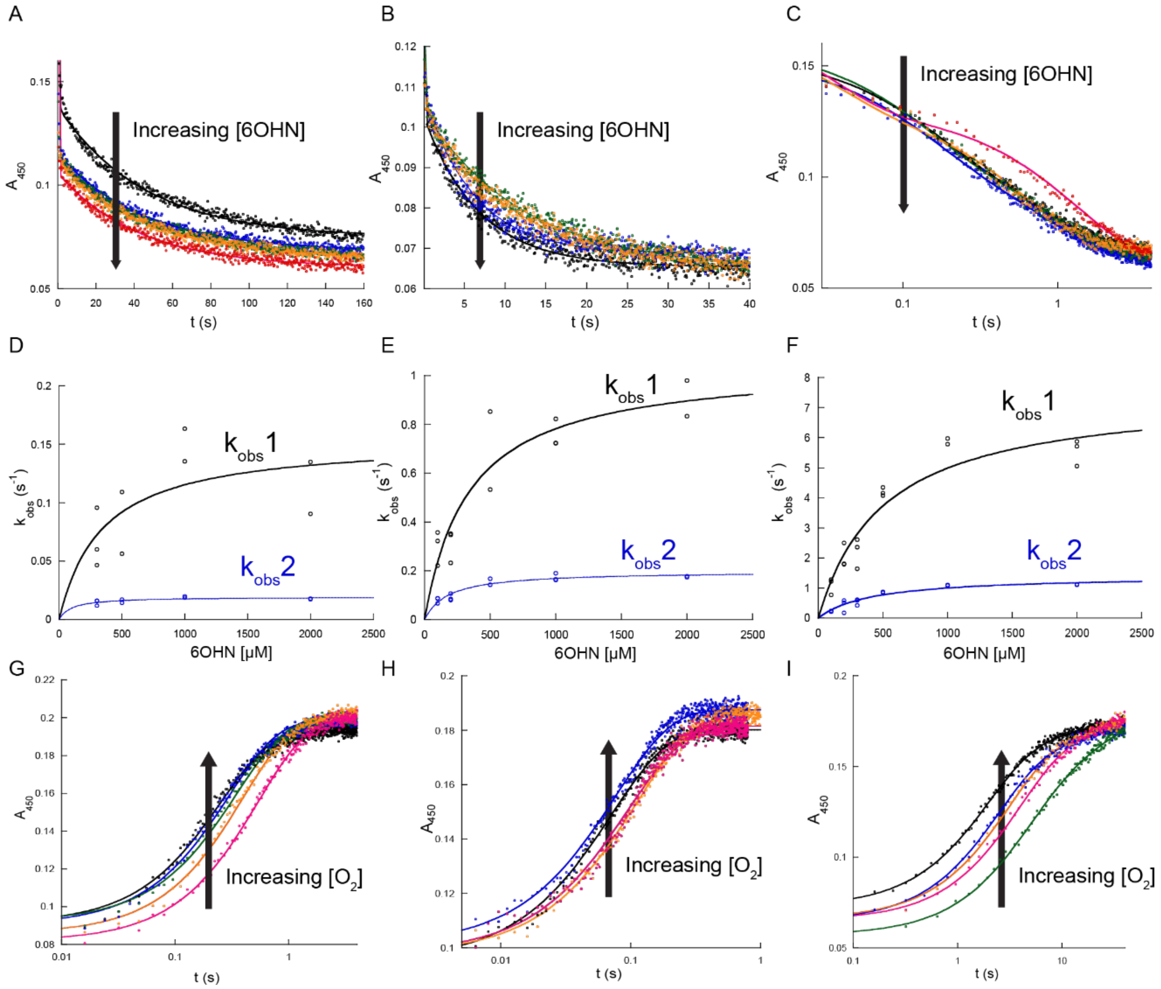


**Fig. S6.** Reaction traces and kinetic analysis of Node D single mutants. (A), (B), and (C), Absorbance trace overlay at 450 nm for the reduction of D→C A90V, M100V, and E300T by various concentrations of 6OHN, respectively. Traces were fitted with equation 2 with two exponential phases. Note the time scale for E300T is logarithmic. (D), (E), and (F), $k_{obs}$ values for the first and second phase plotted against the concentration of 6OHN for D→C A90V, M100V and E300T, respectively. Data for both $k_{obs}$ sets are fitted to equation 4 and parameters from the resulting fitting can be found in Table S1. (H), (I) and (J), Absorbance traces at 450 nm for the oxidation of D→C A90V, M100V, and E300T respectively, containing reduced flavin by various concentrations of $O_2$. Kinetic traces were fit to equation 1 and plots of $k_{obs}$ values against $O_2$ concentration can be found in Fig. 6B.

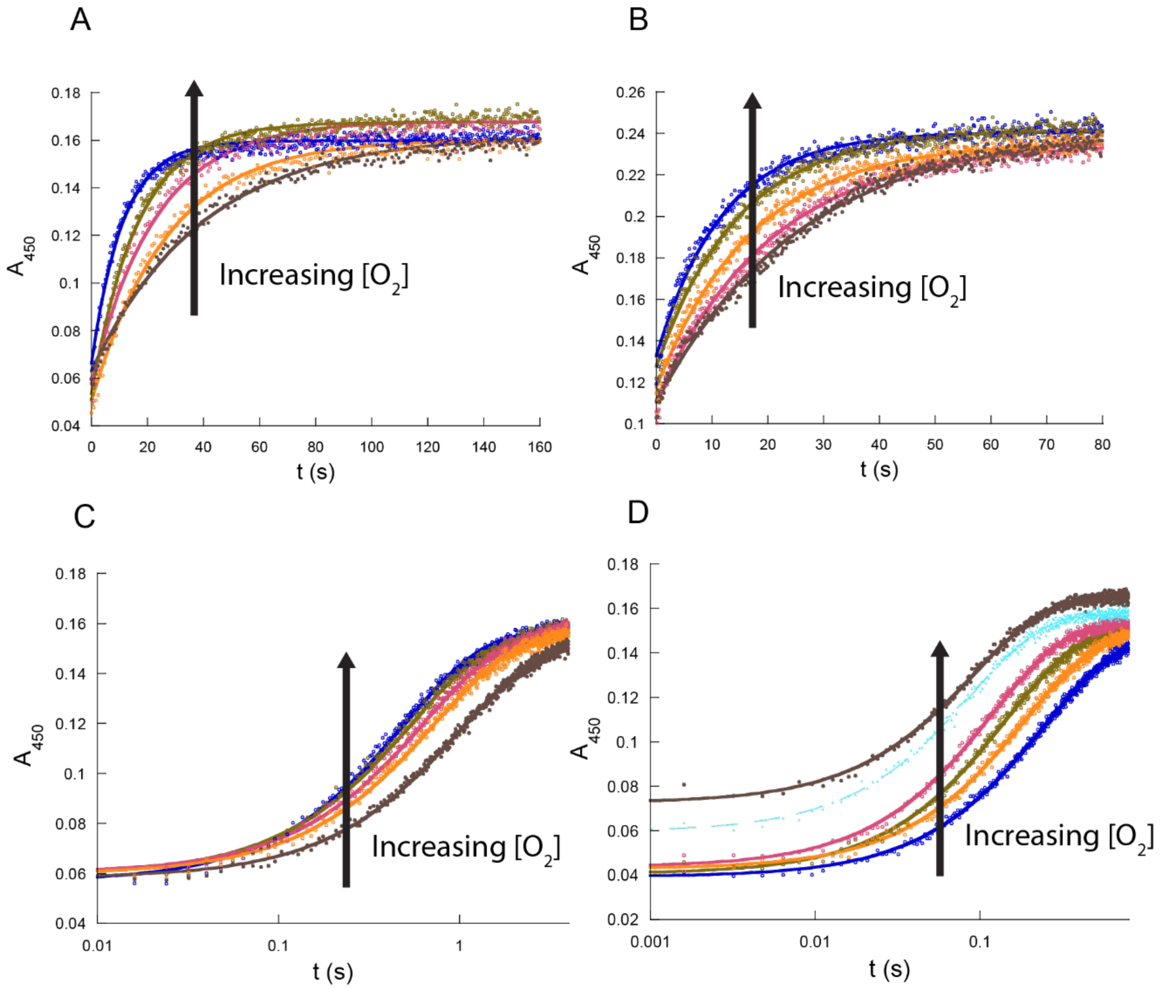


**Fig. S7.** Reaction trace overlay for C→D variants with $O_2$. (A), (B), (C), and (D), Absorbance traces at 450 nm for the oxidation of reduced Node C T300E, Node $C^{5}$, Node $C^{13}$, and Node $C^{21}$, respectively, by various concentrations of $O_2$. Note the logarithmic timescale for panels C and D. Traces were fitted using equation 1.

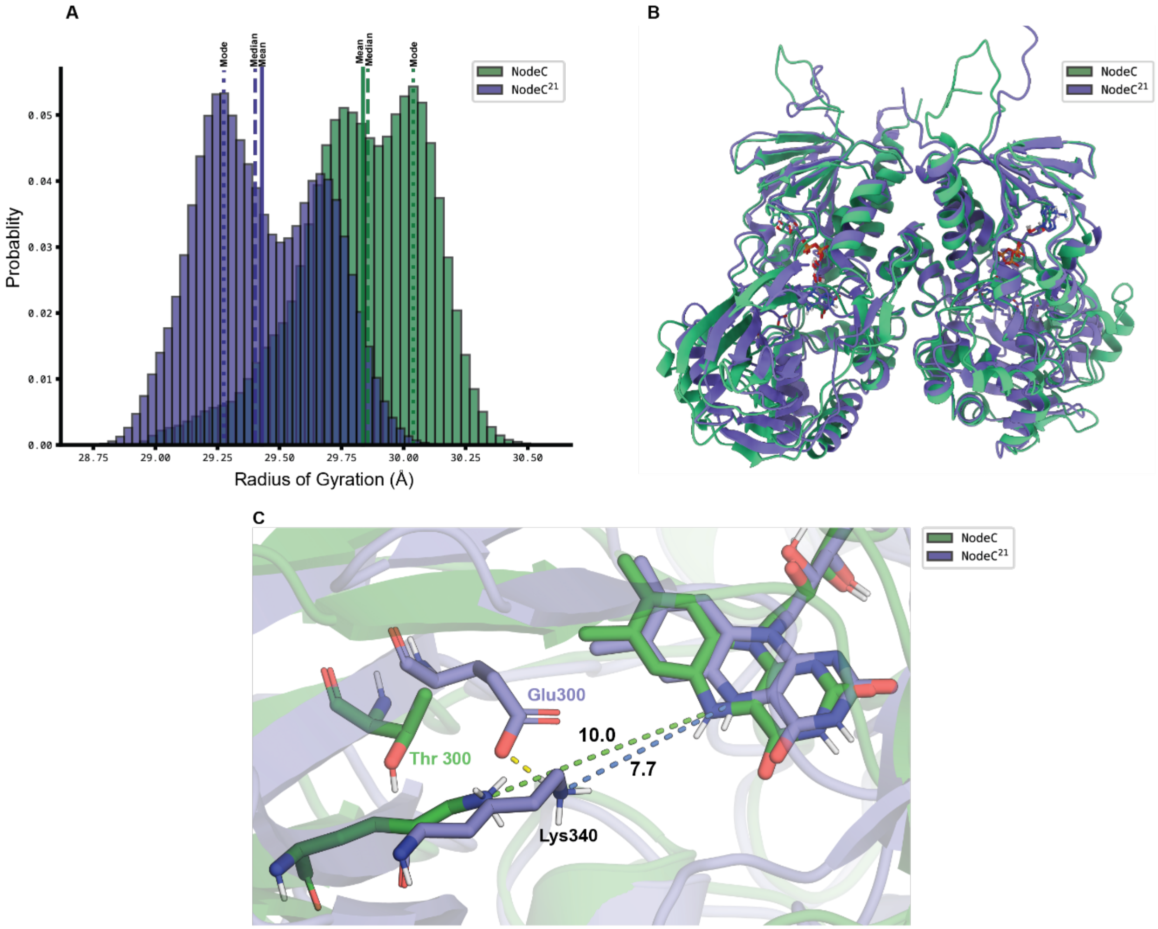


**Fig. S8. Radius of gyration measurements.** (A) Histogram showing the radius of gyration (rgyr) distributions for Node C (green) and Node $C^{21}$ (purple), with vertical dashed lines indicating modal values. $NodeC^{21}$ consistently occupies a smaller volume compared to Node C. (B**)** Representative structures of Node C (green) and Node $C^{21}$ (purple) extracted at the modes of their rgyr distributions, aligned to highlight differences in the volumetric spread of the substrate binding domains. (C) Zoomed-in view of the active site for the structures shown in Fig. S8B. The magnified active site shows the distance between lysine and the flavin isoalloxazine ring in Node C (10.0 Å, green dotted line) and Node $C^{21}$ (7.7 Å, blue dotted line) at this representative snapshot, demonstrating how the overall protein compaction affects the relative position of the lysine.

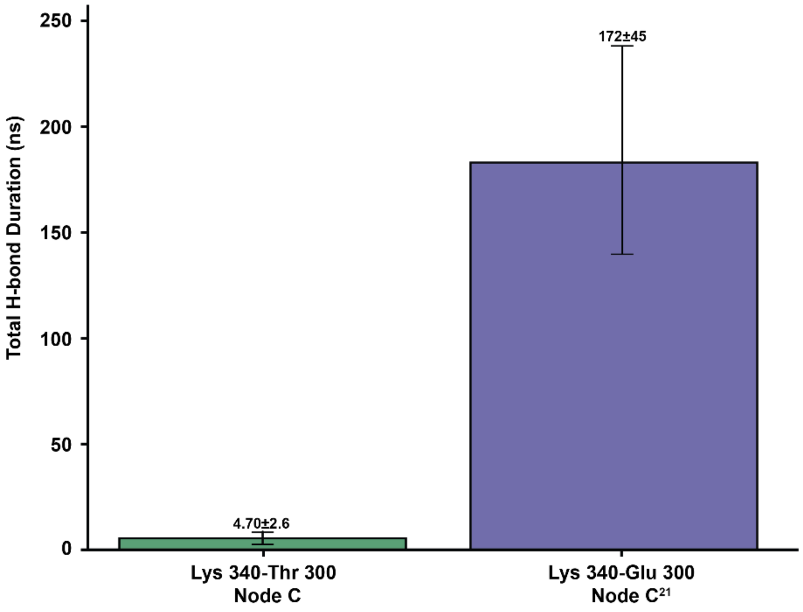


**Fig. S9 -** Hydrogen bond analysis using cpptraj shows a high hydrogen bond contact between Lys340-Glu300 residue pairs in Node C[21] compared to Lys340-Thr300 in Node C.

**Table S1. Kinetic Parameters of NctB, ancestral FAOs and Node D variants[a]**

| | NctB | Node B | Node C | Node D | Node D A90V | Node D M100V | Node D E300T |
|---|---|---|---|---|---|---|---|
| $K_{d1}^{6OHN}$ (μM) | 150±15 | 159±21 | 38±8 | 130±20 | 340±250 | 340±100 | 510±110 |
| $K_{d2}^{6OHN}$ (μM) | 48±7 | 160±12 | 42±29 | 43±5 | 99±50 | 180±40 | 420±80 |
| $k_{red1}^{6OHN}$ ($s^{-1}$) | 180±10 | 2±0.1 | 4.7±0.4 | 12±1 | 0.15±0.04 | 1.1±0.1 | 7.5±0.6 |
| $k_{red2}^{6OHN}$ ($s^{-1}$) | 17±1 | 0.7±0.1 | 1.4±0.4 | 1.1±0.1 | 0.020±0.001 | 0.20±0.01 | 1.4±0.1 |
| $K_{d1}^{Nic}$ (μM) | 540±50 | N.D.[c] | 330±60 | 220±40 | N.D.[e] | N.D.[e] | N.D.[e] |
| $K_{d2}^{Nic}$ (μM) | N.D.[b] | N.D.[c] | N/A[d] | 190±50 | N.D.[e] | N.D.[e] | N.D.[e] |
| $k_{red1}^{Nic}$ ($s^{-1}$) | 0.15±0.01 | N.D.[c] | 0.015±0.001 | 0.053±0.002 | N.D.[e] | N.D.[e] | N.D.[e] |
| $k_{red2}^{Nic}$ ($s^{-1}$) | 0.017±0.001 | N.D.[c] | N/A[d] | 0.014±0.001 | N.D.[e] | N.D.[e] | N.D [e] |
| $k_{ox}^{O2}$ ($M^{-1}s^{-1}$) | 3.8±0.1*10$^4$ | 76±1 | 18±1 | 3.9±0.1*10$^4$ | 7.0±0.1*10$^3$ | 2.8±0.1*10$^4$ | 1.1±0.2*10$^3$ |

[a] N.D. means not determined. Error values denote standard errors.
[b] $K_d$ value is too small to determine from the plot in Fig. S2K
[c] Due to the slow reaction between Node B and nicotine in Fig. S2C, kinetic parameter between Node B and nicotine were not determined.
[d] Only one exponential is observed for the reaction of Node C with nicotine in Fig. S2D
[e] Kinetics of reduction by nicotine were not measured for these variants.

**Table S2** Residue-residue contact frequencies averaged over 3 simulation runs.[1]

| Residue 1 | Residue 2 | Residue Contact Node C (%Avg over 3 runs) | Residue Contact Node $C^{21}$ (%Avg over 3 runs) | Δ%Contact (Node $C^{21}$-Node C) |
|---|---|---|---|---|
| **Lys340** | **Thr300** | **42.55** | | **53.15** |
| **Lys340** | **Glu300** | | **95.70** | |
| **Lys340** | **Glu328** | **40.60** | | **-34.06** |
| **Lys340** | **Tyr328** | | **6.53** | |

[1]Green and purple shaded cells indicate the contact in Node C and Node $C^{21}$ respectively. A contact is defined when any non-hydrogen (heavy) atom from one residue comes within 4.5 Å of any heavy atom in the other residue. This analysis includes the backbone heavy atoms such that there are frames where Lys340 is in contact with both Glu300 and Tyr328 at the same time in Node C21, allowing the total between the two residues to exceed 100%.

**Ancestral protein sequences used in this study**

Node A:
MTKAGHSASGEADYDVIVIGGGFAGVTAARELGLQGYKVLLLEARSRLGGRTYTSKFA
GREVEFGGAWVHWLQPHVWAEMQRYGLGVEEDPLTNLDKALVMLSDGKVKELSPEK
FFHNIREAFEKFCADAREMFPRPYEPMFNPKVKELDKLSVADRIETLDLSEAQRALLNA
MMTLYAGGPTDEFGFASMLKLYACAGWDYDAFMDAETHYRIEGGTMGLIKAMLEDS
GAEVRLSTPVKAVEQESDGVRVTTEDGETITARAVVITVPLNTYKNIEFTPALSPEKQAFI
KEGQMSKGAKIYVHVKQNIGNVFAFCDEPHPLNWVQTEDYGDELGTILSITVARSSLIDI
NDQDAVEKEIRKLFPDVEVVGVSAYDWASDPYSRGAWPAYRVGQLSRYEDMQKPEGR
IFFAGAATANGWHEFIDGAVESGLRAGREVKEFLGLEHHHHHH

Node B:
MTKAGHSASGEADYDVIVIGGGFAGVTAARELGVQGYRVLLLEARPRLGGRTFTGKFQ
GRKVEFGGACVHWIQPHVFAEMQRYGFGFEEVPLANLDKAYVMLSDGKVRDISPEKFD
REYNDAFEKFCARSRELFPRPYSPFFNPEVLELDNVSAAEHIETLGLNEIQRATLNAEMT
LYAGAPTTEFSYTSFVKLFALAAWDYYTFTDSEKHYRIANGGTLGLCKAILDDSGAEVR
LGTVVKAVEQEQDGVRITTADGQTVTAKAVVITVPTKVYPNIEFSPGLSAEKQAFIKNGE
MCDGATMYVRVKQNIGNTFAFCDDPNPFNAIQTEEFDDEIGTILKVTLGRQSLIDINDQD

AVKKEIRRLFPDVEIVGISSYNWATDPFSRQAWPAYRVGWFSKYKDMAKPEGRLFFAG
SATADGWHEYIDGAVESGIRAGREVRELLGREAASRELEHHHHHH

Node C:
MTKAGHSASGEADYDVIVIGAGFAGVTAARELAAQGWRVLVLEARPRIGGRTFTGKFL
GRKIELGGASVHWVQPHVFAEMQRYGFGFEEVPLANLDKAYVMLSDGKVHDVPPEKF
DREYNDAFDKFCARSRELFPRPYSPFANPEVLKLDNISAAEHIETLGLNEIQRASLNAELT
LYAGAPTTEFSYTSFVKLFALAAWDTYTFTDSEKHYHIANGGTAALCKAILDDSKADVR
LGTVVKAVEQSEDGVTLTTADGQTVTAKAAVLTLPTKVYPDIEFKPGLSAEKQAFIKNG
EMCDGATMYVRVKQNLGNTFAFCDDPNPFNAIQTEEFDDEIGTILKVTLGRQSLIDINDQ
DAVAAEIRKIHPDVEIVDIAPYNWAKDPFSKQAWPAYRAGWFSKYKDMAKPEGRLFFA
GSATADGWHEYIDGAVESGIRAGREVRQLLEKENASLEHHHHHH

Node D:
MTKAGHSASGEADYDAIVVGAGFAGVVAARELAAQGRRVLVLEARDRIGGRTFVGTFL
GRRIELGGAGVHWVQPHVFAEMQRYGFGFKEAPLANLDKAYMMLSDGRVLDVPPDTF
DREYNAAFDKFCARSRELFPRPYSPLDNPEVLALDGVSAHEHLESLGLNELQRASMNAE
LTLYGGAPTTELSYPSFVKFHALASWDTYTFTDSEKRYHVAGGTHALCEAILDDARADV
RLGAPVEAVAQGENGVTLTLADGQTFSASTAVLTLPTKVYPDIAFEPPLSAEKQAFIENA
EMCDGAELYVHVRQNLGNTFAFCDDPNPFNAIQTYAYNDELGTILKITLGRQSLIDIEDI
DAVAAEIRKIHHDVEVLSIAPYNWAKDRFSKTAWPAYRKGWFSKYKDMAKPEGRLFF
AGSATADGWHEYIDGAVESGIRAGREVRRFLENPGASLEHHHHHH
